# Anatomy of the complete mouse eye vasculature in development and pathology explored by light-sheet fluorescence microscopy

**DOI:** 10.1101/2022.12.20.521194

**Authors:** Luc Krimpenfort, Maria Garcia-Collado, Tom van Leeuwen, Filippo Locri, Anna-Liisa Luik, Antonio Queiro-Palou, Shigeaki Kanatani, Helder André, Per Uhlén, Lars Jakobsson

## Abstract

Eye development and function rely on precise establishment, regression and maintenance of its many sub-vasculatures. These crucial vascular properties have been extensively investigated in eye development and disease utilizing genetic and experimental mouse models. However, due to technical limitations, individual studies have often restricted their focus to one specific sub-vasculature. Here, we apply a workflow that allows for visualisation of complete vasculatures of mouse eyes of various developmental stages. Through tissue depigmentation, immunostaining, clearing and light-sheet fluorescence microscopy (LSFM) entire vasculatures of the retina, vitreous (hyaloids) and uvea were simultaneously imaged at high resolution. In silico dissection provided detailed information on their 3D architecture and interconnections. By this method we describe remodelling of the postnatal iris vasculature following its disconnection to the feeding hyaloid vasculature. In addition, we demonstrate examples of conventional and LSFM-mediated analysis of choroidal neovascularisation after laser-induced wounding, showing added value of the presented workflow in analysis of modelled eye disease. These advancements in visualisation and analysis of the respective eye vasculatures in development and complex eye disease open for novel observations of their functional interplay at a whole-organ level.

## Introduction

Ocular vascular alterations are frequent causes of visual impairment. Defects in vascular permeability resulting in oedema and bleeding, as a consequence of pericyte loss and basement membrane disruption, are common denominators. In addition, mispatterning through neovascularization and microvascular tuft formation can be detrimental. Hyaloids instead, mainly cause disease upon postnatal pathological persistence. These pathological features are often localised to specific regions or sub compartments of the eye and to stages of vascular development, characterised by extensive vascular morphogenesis. Targeting these pathogenic properties specifically, while preserving crucial normal function, has provided challenges. Nevertheless, inhibition of angiogenesis and permeability by administration of blocking antibodies to Vascular endothelial growth factor A (VEGFA), comprises standard treatment for several ocular pathologies, including age-related macular degeneration (AMD) and diabetic retinopathy^1^. However, side effects, gradual treatment resistance and relapse are common, illustrating the need for deeper knowledge on the heterogeneity between and within the sub compartments and vasculatures of the eye.

Genetic and experimental mouse models have provided fundamental knowledge on mechanisms of vascular morphogenesis, malformation and malfunction related to eye disease. Individual studies have traditionally restricted their scope to one specific vascular bed of the eye, potentially overlooking key phenotypes in any of the other interconnected vascular compartments (i.e. retinal vasculature, hyaloids, neuronal retina, choroids, lens, limbus, iris)^2^ crucial for eye development and function. In the adult mouse eye, the choroid and the dense fenestrated choriocapillaris support the retinal pigment epithelium (RPE) and the outer neuronal retina^3^. The inner retina is supplied by three layers of the retinal vasculature, comprising the blood-retinal barrier through which transport is highly restricted. The iris and cornea are respectively supported by the iris- and the limbal vasculature. In embryonic development, the lens is supported by the transient hyaloid vasculature composed of the hyaloid artery (HA), the vasa hyaloidea propria (VHP), tunica vasculosa lentis (TVL), and the most anterior pupillary membrane (PM) that drain via the iris into the choroid venules. As the eye develops and the inner neuronal retina becomes vascularised and oxygenated, the hyaloids regress, leading to dramatic alterations in flow patterns. In the mouse, the first signs of regression appear around postnatal day 4 (P4) and is completed sometime after P20, though it is unclear exactly when^4–6^. Hyaloid regression is a prerequisite for full vision, as evident in the congenital disease persistent foetal vasculature (PFV), that may cause childhood blindness^7^.

The vast majority of studies in experimental mouse models of vascular eye disease have relied on dissection of retinas or other sub compartments followed by immuno-labelling, flat-mounting, microscopy and image analysis^8^. These procedures rarely allow for preservation of all eye compartments and hence limit data collection. In recent years, optical clearing of tissues and organs have developed to a point where practically any type of tissue can be made transparent and compatible with Light-Sheet Fluorescence Microscopy (LSFM). However, the mouse eye holds challenges for optical clearing and immunolabelling, due to light absorption by melanin and poor antibody penetration of the sclera. To overcome this, current methods apply intensive bleaching at high temperatures^9–11^, antibody injections, complete resection of the sclera and the RPE^12,13^, or use of enzymes to permeabilise the sclera^11^. Albino mice requires less extensive depigmentation, making them attractive for LSFM as demonstrated by Darche et al^5^. However, genetically modified strains on C57Bl6 background are dominating, for which this step is crucial.

Here we describe a workflow of optical tissue clearing, antibody labelling and LSFM imaging that together with in silico dissection permit visualisation and analysis of the complete mouse eye vasculature in 3D. From a single mouse eye, extensive information on the uveal, retinal and hyaloid vasculatures can be retrieved. The whole-eye perspective further enables quantitative analysis of the asymmetrical development of the retinal vasculature, as well as the postnatal development of the hyaloid and iris vasculatures. Analysis of mouse eyes following laser-induced choroidal neovascularisation (CNV) exemplifies how this methodology can enhance contextual information in models of complex eye disease. The workflow hence comprises an easy and robust pipeline for analysis of the complete mouse eye vasculature in development and disease models.

## Materials and Methods

### Animals

Experiments included male and female mice from pure C57Bl6/J or mixed with 129S6. Eyes were taken from mice at P2 through adulthood, fixed in 4% PFA for 4h at 4°C and stored in phosphate-buffered saline (PBS) at 4°C until further use. Animal experimental protocols were approved by the Stockholm North Ethical Committee on Animal Research (permit numbers 9780-2019 and 7053-2020.) All animal experiments were carried out in accordance with their guidelines as well as in accordance with the ARVO statement for the Use of Animals in Ophthalmologic and Vision Research.

### Laser-induced wounding for choroidal neovascularisation

Eight-weeks-old C57BI6/J mice (Charles River, Cologne, Germany) were anesthetized by inhaled isoflurane in room-air. Both pupils were dilated by topical administration of tropicamide (0.5%). Choroidal neovascularisation lesions were induced in both eyes by a 532 nm green laser photocoagulator (Phoenix-Micron Inc.) set for 50 μm spot, 180 mW intensity, 100 ms duration, with 3 shots per eye. After laser-induction, animals were hydrated with 500 μL of saline (9 mg/mL) subcutaneously while eyes were kept lubricated by topical administration of Viscotears. Seven days post laser induction, mice were anesthetized by isoflurane and their pupils dilated by tropicamide eyedrops. Spectral domain optical coherence tomography (SD-OCT) was performed with the assistance of a live-fundus image (Phoenix-Micron Inc). B-scan were acquired midline to the lesion centre by averaging 50 photograms. Fluorescence fundus angiography was performed after the acquisition of OCT B-scans by subcutaneous injection of 30 mg/kg sterile fluorescein and imaged with fixed camera exposure settings 5 min after injection.

### Materials and solutions

*Solution-1* consists of 8 to 10% (v/v) THEED, 5% (v/v) Triton X-100, and 25% (w/v) urea in dH_2_O. PTx.2 consists of 1x PBS with 0.2% Triton X-100. Permeabilization solution consists of PTx.2 with 2.3% glycine (w/v) and 20% DMSO. Blocking solution consists of PTx.2 with 6% donkey serum (Jackson IR, AB_2337254) and 10% DMSO. PTwH consists of 1x PBS with 0.2% Tween-20 and 0.1% of 10mg/mL heparin (vol/vol).

### Depigmentation

After fixation, mouse eyes were washed with PBS and exposed to a combined depigmentation and bleaching protocol. This protocol combines the methods of CUBIC-based DEEP-Clear^14^, ECi-based EyeCi^9^ and iDISCO+ based EyeDISCO^15^. In brief, eyes were exposed to pre-chilled acetone (−20°C) for 2h and washed with PBS 3 x 2 mL at room temperature (RT), followed by depigmentation by 3h incubation in *Solution-1* (37°C, gentle shaking) and 3 x 2 mL PBS wash at RT for 5 min. Then, the eyes were bleached with 10% H2O2 in PBS at 37°C (gentle shaking) until pigmentation was lost, ranging from 1h to overnight, for P2 eyes and adult eyes, respectively.

### Immunostaining of whole eyes

Following H2O2-bleaching, the eyes were immuno-stained according to the iDISCO+ method (https://idisco.info/idisco-protocol/, December 2016)^16^. In short, eyes were washed twice in PTx.2 for 1h at RT, placed in permeabilization solution for 24h at 37°C, then blocking solution for 24h at 37°C. This was followed by primary Ab incubation in PTwH/5% DMSO/3% donkey serum at 37°C for >36h after which the eyes were washed 4 – 5 times over 24h in PTwH, at RT. The eyes were incubated with secondary Ab in PTwH/3% donkey serum at 37°C for at least 36h and washed 4 – 5 times over 24h with PTwH at RT. Secondary antibodies were spun down at 16100 g for 2 minutes prior to use.

### Clearing

The bleached eyes were embedded in gelatin (3% in PBS, Uhlén lab, unpublished) in 35 mm cell culture dishes, cooled to 4°C and cut out into blocks, further stored in PBS. The embedded eyes were cleared and refractive index (RI)-matched according to a slightly altered iDISCO+-based EyeDISCO protocol^16^, summarized as follows. The eyes were dehydrated in methanol/dH_2_O-series (20%, 40%, 60%, 80%, 100%, 100%), 2h per step, at RT with gentle shaking. Eyes were then placed in 67% dichloromethane (DCM) and 33% methanol for 3h at RT with gentle shaking, followed by two washes in 100% DCM for 15 min each, at RT with gentle shaking. Finally, eyes were RI-matched by incubating for at least 5h in DiBenzyl Ether (DBE) at RT, though preferably overnight with a change of DBE after 4 hours. After the RI-matching, eyes were stored in the dark at RT or imaged.

### Relabelling and flat mounting of retinas and cleared eyes

After imaging, eyes were retrieved from gelatin for subsequent re-immunostaining and flat mounting. Eyes in gelatin first need to be rehydrated using the reverse methanol/dH_2_O series (100%, 100%, 80%, 60%, 40%, 20%), 1h per step at RT with gentle shaking/rocking. (Note: any remaining DBE in the eye will react with the water in the rehydration steps to form white crystals, this does not render the eyes unusable, but should ideally be prevented by an extra incubation in 100% methanol). After rehydration, eyes were cut out of the gelatin and subsequently dissected. Then eyes were incubated in blocking/permeabilization solution (5% donkey serum (v/v), 1% BSA (w/v), 0.5% Triton X-100 (v/v) and 0.01% NaN_3_ in PBS) for 1h at RT with gentle rocking. Immunolabelling was done by overnight incubation with primary antibodies in 1:1 PBS:blocking/permeabilization solution, followed by washing 3x in PBS (0.01% Triton X-100) for 15 minutes. The same was done for secondary antibodies whereafter dissected eye compartments were flat mounted with addition of the mounting media Prolong Gold.

### Imaging

The cleared eyes were imaged in the LaVision Biotec UltraMicroscope II, using the Olympus MVX10 zoom body, internal 2x objective (immersion lens) and Andor Zyla 5.5 sCMOS. For whole-eye acquisitions, lens zoom was set to 2x (effectively 4x), z-step to 1.93 μm, light-sheet thickness to 3.86 μm, light-sheet width to 50%, laser power was set to fill the dynamic range of the camera, and frame acquisition time was 100 ms. Samples were excited by three light-sheets from the left, and dynamic horizontal focus over the whole eye was used in all cases. Chromatic aberration correction was set to 0 for all channels and focused optimized for the CD31 channel. For higher-resolution imaging, zoom was set to 6.3x (effectively 12.6x), light-sheet width was set to 10%, objects were lighted with a single left-sided laser and chromatic aberration correction was applied. All other settings were left identical.

### Image analysis

The raw 3D image stacks (.tif format) were loaded into ImageJ (version 2.9), merged into one TIFF per acquisition and cropped to the relevant area. If necessary, different colour channels were aligned first. TIFFs were loaded into Imaris (Oxford Industries, versions 9.7.2 – 9.9.1), for further analysis. The oblique slicer tool was used to make cross sections of varying thickness in any orientation required. For in silico dissection the surface tool was used manually to highlight desired regions. The in silico dissection process is described step-by-step in the supplemented beginners tutorial. For the analysis of the retina, the choroid arteries were first highlighted using the measurement point tool (Suppl. Fig. 3A), so the orientation of the sample was easily determined. Following this, the oblique slicer tool was used to create a 100 μm section, on the surface of which a measurement point line was drawn to acquire the radial expansion of the superficial and deep layers in 4 directions (Suppl. Fig. 3A). Next, the whole retina was dissected in silico, followed by dissection of a smaller slice in the dorsal region of the eye, which includes one vein and one artery (Suppl. Fig. 3B). This region was dissected into the superficial, intermediate and deep layer, of which surface volumes were subsequently created. Finally, ERG nuclei (Suppl. Fig. 3C) were counted using the spots tool. Spot quality, volume and distance to vessel layer surfaces were used (in indicated order) to filter out antibody precipitates and noise. Settings were calibrated by expert supervision and kept identical within each eye.

For experimental details and a step-by-step protocol, see Supplementary material and methods.

## Results

### Bleaching, immunolabelling and clearing of intact mouse eyes enable 3D visualisation and in silico dissection of all ocular vasculatures

Application of LSFM to image antibody-labelled structures of the intact mouse eye, requires complete antibody penetration and tissue transparency. In the eye, the sclera comprises a rigid structure known to obstruct antibody penetration. Furthermore, RPE cells contain melanin that absorbs light of various wavelengths, thereby restricting excitation and emission of structures of interest. To increase antibody penetration and reduce the impact of melanin while preserving tissue architecture, we combined and adjusted steps of the protocols of EyeCi^9^, EyeDISCO^15^ and DEEPClear^14^. In brief, excised eyes with optic nerves were fixed in 4% PFA in PBS (Fig. 1A), submersed in 100% acetone, depigmented in solution 1 (see methods) and bleached in 10% hydrogen peroxide at 37°C (Fig. 1B). For immunolabelling and RI tissue-matching to permit transparency, a modified iDISCO+ protocol was applied as follows. The sample was permeabilized in a 20% DMSO solution and blocked for unspecific binding by donkey-serum. Incubation with primary-, followed by secondary-antibodies, embedding in 3% gelatin and dehydration in a step-wise low-to-high methanol series prepared for the final steps of clearing. DCM and subsequently DBE were used to adjust the RI to that of the tissue (clearing), making the eyes transparent (Fig. 1C). For detailed experimental protocol, see Supplementary methods. The implemented changes from previously published protocols include incubation of eyes in Solution-1 before, instead of after, the hydrogen peroxide-solution as well as lowering the temperature for hydrogen peroxide incubation from 55°C to 37°C. This eliminated bubble formation and reduced damaging effects on tissue integrity while preserving bleaching efficiency (data not shown).

**Figure 1.**
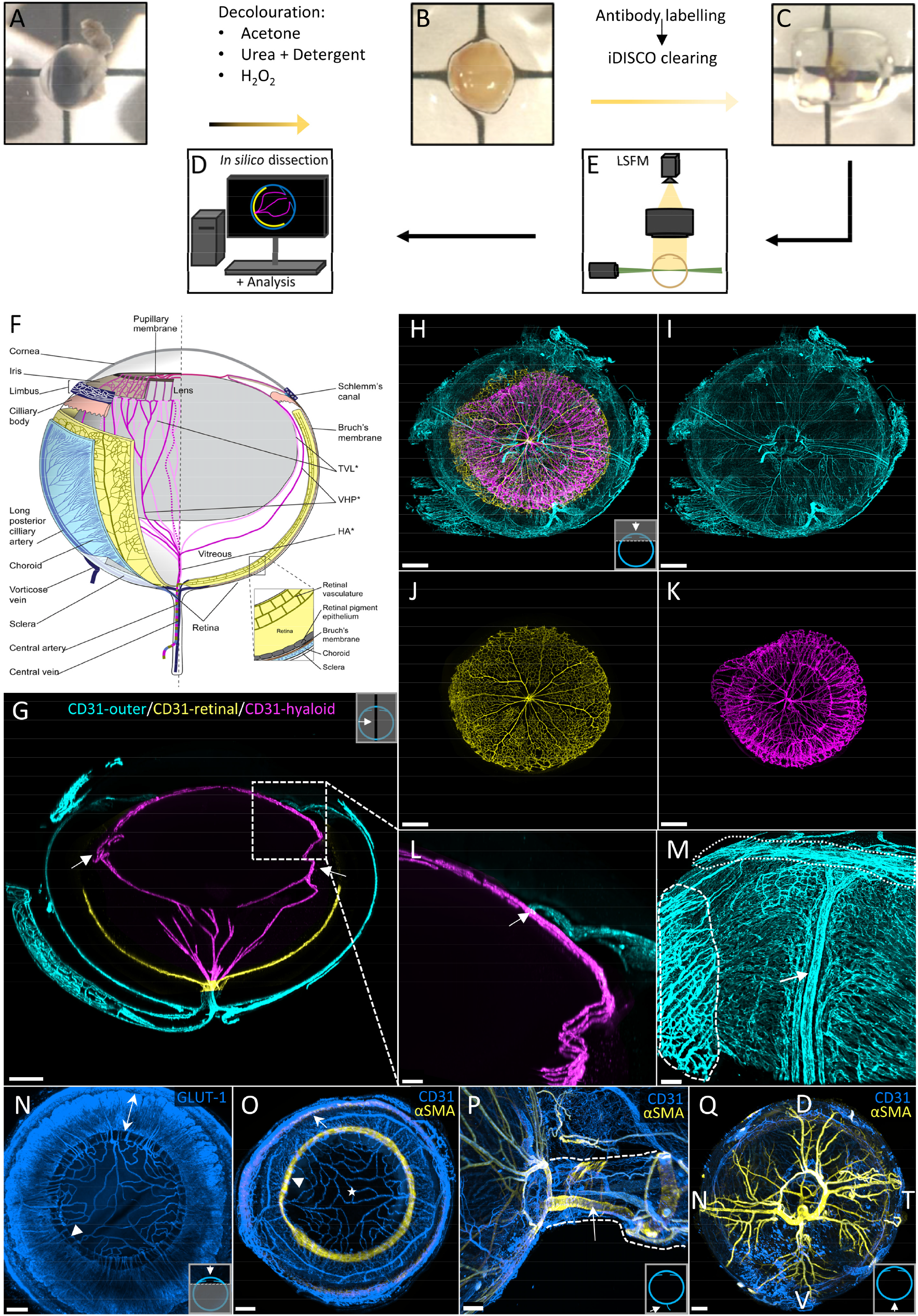
Optical clearing combined with LSFM allows for visualisation of the entire eye vasculature. (**A**) A freshly harvested mouse eye after fixation, (**B**) decoloured using acetone, solution 1 (urea, THEED and Triton-X100) and 10% hydrogen peroxide and (**C**) subsequently immunolabelled, embedded in gelatin and made optically transparent with the iDISCO+ clearing protocol. (**D**) Schematic outline of the LSFM setup and, (**E**) of computer-assisted image analysis. (**F**) Schematic representation of the whole-eye and its vasculatures at P10. TVL, tunica vasculosa lentis; VHP, vasculosa hyaloidea propria; HA, hyaloid artery. (**G-M**) In silico dissected CD31+ structures of a whole P6 eye pseudo coloured to highlight the outer vasculature (including muscle and choroids, cyan), retinal vasculature (yellow) and hyaloid vasculature (magenta). (**G**) A 200 μm optical cross section of the eye (arrow in inserted box, upper right corner indicates direction of presented view, where the black area is presented, grey is not). Arrows indicate vessel artefacts due to lens expansion during preparation. (**H-K**), anterior perspective of the same eye, the iris/most anterior part of the eye is omitted for clarity. (**I**) The outer vasculature. (**J**) The retinal vasculature. (**K**) The hyaloid vasculatures. (**L**) Arrow points to direct connections between the hyaloid vasculature and the iris vasculature. (**M**) Higher magnification of the choriocapillaris, also showing vessels of the eye muscles (left, dashed line) and limbus (up, dotted line). The arrow indicates the ciliary artery feeding the limbus, iris and ciliary body. (**N**) Anterior view of the iris (double arrow) and PM (P5), labelled for GLUT-1. Note absence of signal in the pupillary muscle (arrowhead). (**O**) Anterior view of the anterior part of the eye at P4, labelled for CD31 (blue) and alpha smooth muscle actin (αSMA, yellow). αSMA indicates the ciliary muscle at the base of the iris (arrow) as well as the pupil muscle (arrowhead), surrounding the PM vasculature (star). (**P**) Close up of the optic nerve entering the adult eye (dotted lines), arrow indicates central retinal artery. CD31 (blue), αSMA (yellow). (**Q**) Posterior perspective of same adult eye; the large choroid arteries can be used to orient the eye. V=ventral, D=dorsal, T=temporal, N=nasal. Labelling: CD31 unless otherwise specified. Scale bars: H-K, M-O, Q = 200 μm, G, P = 150 μm, L = 50 μm.

Application of CD31 antibodies within the above protocol, followed by LSFM (Fig. 1D), allowed for acquisition of high-resolution images (up to 0.48 μm/pixel in xy, Z-step size of 1.93 μm) of all vasculatures and their connections throughout the entire eye (Fig. 1F-K). To permit analysis and visualisation of individual vascular beds, data (1200-1800 z-planes/eye) was imported to the image software Imaris and processed in 3D. Based on their specific spatial localisation within the eye, the vasculatures were in silico dissected and pseudo-coloured (Fig. 1E, G-K, Sup. Movie #1, described in supplemental tutorial). The choroid and the choriocapillaris (Fig. 1M, Suppl. Fig. 1A&B), as well as the vorticose veins (Suppl. Fig. 1C&D, arrows), the limbus (Suppl. Fig. 1E&F), ciliary body and muscles (Suppl. Fig. 1G-J), Schlemm’s canal (Suppl. Fig. 1I-K) and the lymphatic vasculature (Suppl. Fig. 1J-L), were resolved in detail. Comparison of P4 and adult choroid, vorticose and limbus vasculatures, indicates the ability to follow the extensive development of this compartment (Suppl. Fig. 1A-F). At the same time, details on the retinal vasculature including arteries, veins and capillaries were recorded (Fig. 1J). In silico 3D isolation and colour coding of the retinal and hyaloid vasculatures (Suppl. Movie #2) exposed their respective anatomy and drainage (flow direction inferred by anatomy). Both the retina and the hyaloid vasculature are supplied by branches of the central artery but while the retinal blood drains through the central vein within the optic nerve sheath, the hyaloid vasculature (Fig. 1K) drains into the iris vasculature (Fig. 1L, N&O).

Staining for alpha smooth muscle actin (αSMA) exposed smooth muscle cell-covered vessels of the choroid. These vessels could in turn be used as guides for temporal-nasal and ventral-dorsal anatomical orientation (Fig. 1P, Q)^9^. In addition, the pupil and ciliary muscle stood out as highly positive for αSMA (Fig. 1N, dotted circles). Unfortunately, the clearing procedure frequently damaged the lens that occasionally caused breakage of TVL vessels (Fig. 1G, arrowheads). The results described above demonstrate that immunolabelling, tissue clearing, LSFM and in silico dissection, enable detailed visualisation and analysis of all vasculatures of the mouse eye.

### Relabelling and flat-mounting of previously cleared eyes

LSFM is an excellent technique for acquiring overviews of large samples in 3D with relatively high resolution. However, laser scanning confocal microscopy of flat-mounted samples generally provides images at even higher resolution. To apply these microscopy techniques sequentially on the same sample would permit anatomical 3D analysis from LSFM, and deeper analysis of selected regions by laser scanning confocal microscopy. To test this workflow, eyes previously imaged by LSFM were rehydrated after which gelatin was trimmed away. Following dissection of sub-compartments (retinas and RPE/choroids etc.) samples were re-incubated with antibodies to restore signals after partial fading. Samples were then flat-mounted and imaged by laser scanning confocal microscopy. In this way, retinal vessels and bifurcations previously visualized in 3D by LSFM (Fig. 2A), were identified in flat-mounted samples at high resolution (Fig. 2B). Aside from the retina, also the hyaloid vasculature, choroid vasculature, Schlemm’s canal and limbus vasculature, could be flat-mounted and imaged at higher resolution (Suppl. Fig. 2A-D). This workflow thereby allows for the study of individual compartments and their interactions in the intact eye, followed by high-resolution analysis of dissected regions of interest.

**Figure 2.**
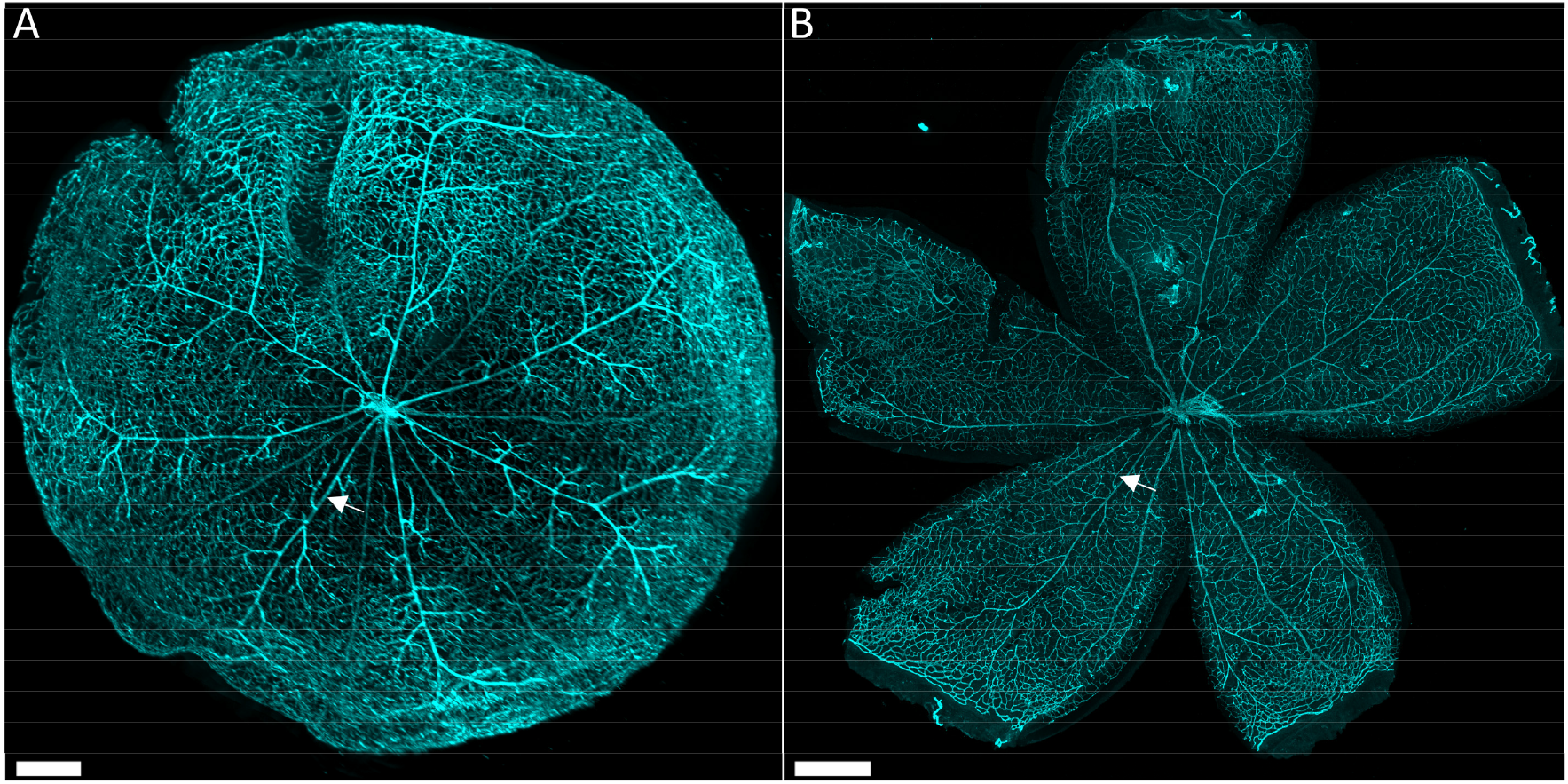
Cleared mouse eye samples can be retrieved for re-staining, flat-mounting and imaging at higher resolution. (**A**) in silico dissected, CD31-labelled P12 retina of a whole cleared eye. (**B**) The same retina after rehydration, physical dissection, relabelling (CD31), flat-mounting and confocal imaging. Arrows indicate identical positions. Scale bars: A = 200 μm, B = 500 μm.

### Analysis of the developing retinal vascular layers in intact eyes exposes asymmetrical expansion

The mouse retinal vasculature starts to populate the retina at birth through outwards radial growth of the superficial layer from the optic nerve. Around P7, just before the superficial vascular plexus has reached the end of the retina, endothelial cells (ECs) start to sprout perpendicular into the neuronal compartment to establish the deep vascular plexus. Shortly thereafter, around P12, the intermediate layer starts to form in between the superficial and deep plexus^8^. The speed by which the individual plexi form, as well as their specific morphologies, are differentially affected by individual signalling pathways (Vascular endothelial growth factor A, WNTs, Transforming growth factor beta)^17,18^. These properties can thereby indirectly reveal alterations in specific signalling cascades. Detailed analysis of the deep and intermediate layers relies on maintained 3D structural integrity, a property that is generally kept in the LSFM protocol but often problematic in 2D flat-mounting due to pressure exerted from the cover slip. In silico dissection of P8, P10 and P12 retinal vasculatures, from whole-eye data acquired by LSFM, allowed for clear visualisation of the retina in 3D (Fig. 3A-C). This enabled examination of progression of the deep layers over time, with initiation below the “oldest” vessels close to the optic nerve followed by expansion towards the periphery (Fig. 3D-F, J, Suppl. Fig. 3A). Further in silico dissection allowed for separation of the three plexi (Fig. 3G-I). Subsequent staining for the EC transcription factor ERG permitted counting of the number of EC nuclei across the different layers over time, albeit subjective to regional alteration in resolution and signal intensity (Fig. 3K, Suppl. Fig. 3C-D). At P8, around 90% of retinal ECs were found in the superficial plexus, but as ECs progressively invade and populate deeper layers, the superficial EC proportion was only 40% at P12. Next, the main choroid vessels were employed to identify the precise anatomical orientation of the eye and hence the retina (Fig. 1Q) as described by Henning et al^9^. This revealed the slowest and fastest progression to occur in the dorsal and ventral sides, respectively (Fig. 3L, Suppl. Fig. 3E-H). Thus, anatomical positioning should ideally be considered when comparing phenotypes relating to invasion and establishment of the different layers. Hereby we exemplified that LSFM of intact eyes can provide detailed data on retinal vascular expansion of all three layers integrated in the whole-eye context.

**Figure 3.**
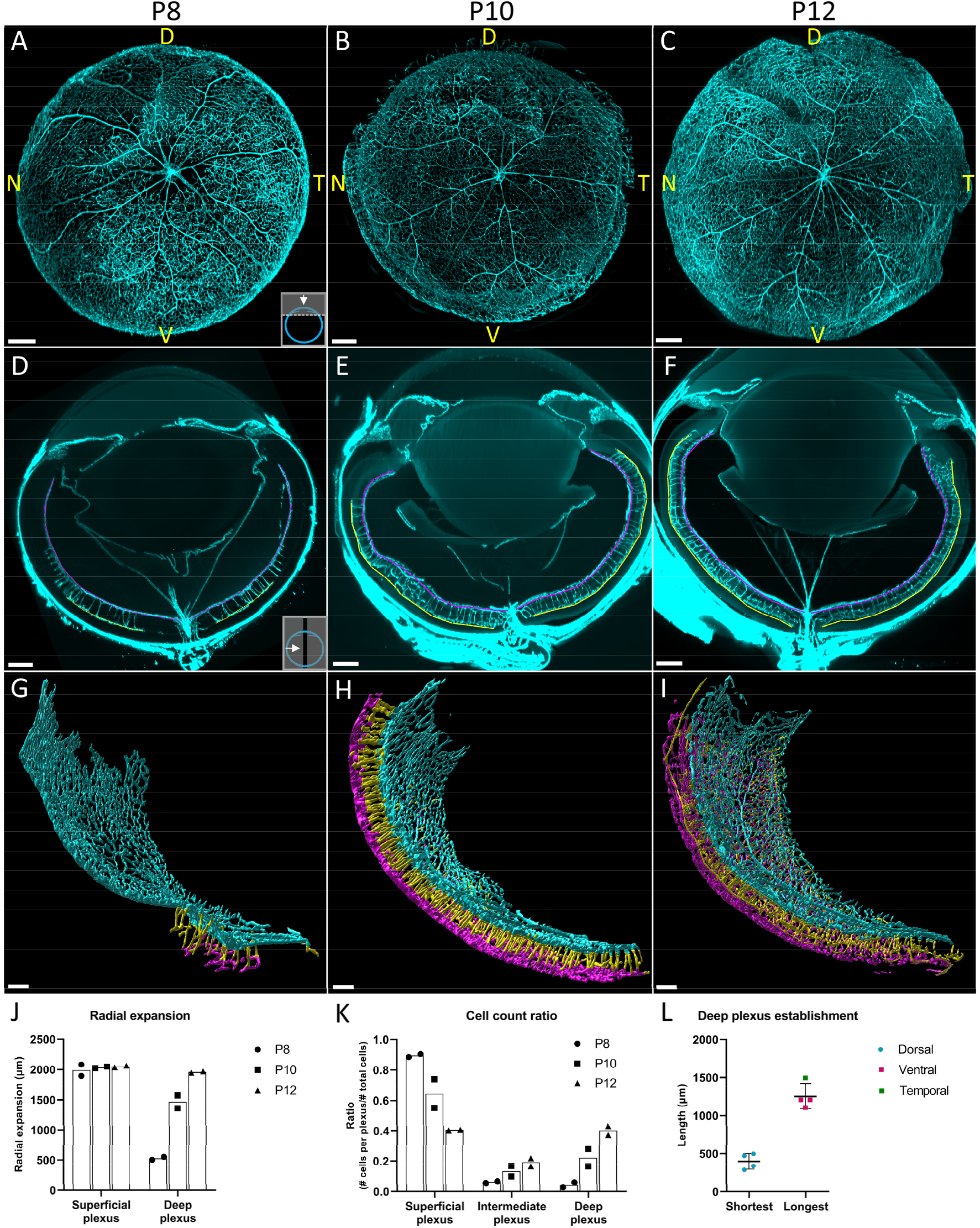
Quantification of retinal vascular development in 3D exposes asymmetrical expansion. (**A-C**) In silico dissected retinas of whole mouse eyes at P8, P10 and P12. V, ventral; D, dorsal; T, temporal; N, nasal. (**D-F**) 100 μm in silico cross sections of P8, P10 and P12 eyes. Magenta and yellow lines indicate measurements of superficial and deep layer establishment. (**G-I**) Sectors of retinas at P8, P10 and P12, in silico dissected to highlight the three vascular plexi individually using surface renderings. (**J**) Quantification of radial expansion of the superficial and deep plexus at P8, P10 and P12. (**K**) Quantification of endothelial cell count ratio (cells in layer/total number of cells) per retinal plexus from P8, P10 and P12. (**L**) Quantification of shortest and farthest deep layer establishment at P8, n=4 retinas. Colour code indicates the direction of the shortest and longest establishments. Labelling: CD31. Scale bars: A-F = 200 μm, G-I = 100 μm.

### Regression of the hyaloid vasculature

In the mouse, the embryonic hyaloid vessels start to regress around P4, a process suggested to be triggered by a decrease in VEGFR2 signalling in hyaloid ECs and promoted by apoptosis signals mediated by perivascular mural cells and macrophages^19–23^. Although kinetics of regression has been characterised, there are discrepancies in literature, potentially related to the fragile nature of free-floating dissociating vessels in the vitreous and the difficulty to analyse them by conventional techniques (e.g. sectioning, dissection, OCT and stereomicroscopy)^4,6,24,25^. A recent report, utilizing albino eyes and LSFM, demark the power of this strategy to assess hyaloid biology^5^. To clarify kinetics of hyaloid regression we applied LSFM on cleared intact eyes of litter mates of P2, P4, P6, P8, P10, P12 and P16 mice. The CD31+ hyaloid vasculatures were in silico dissected based on their spatial localisation and presented from the side (Fig. 4A-G) and top, excluding either the posterior (Fig. 4H-N) or the anterior hyaloid system (Fig. 4O-U). In line with published data, the first signs of hyaloid regression appeared at P4 as solitary vessel fragments^26^ (Fig. 4I, arrow), also pointed out in P6 and P8 (Fig. 4Q, R and X). Vessels of the PM disappeared between P10 and P12, while the VHP and TVL were clearly reduced at P16 in agreement with literature^4,5^. While allowing for in depth characterisation, the protocol caused expansion and rupturing of the lens which in turn led to breakage of the TVL from P2 (Fig. 4Z, magenta arrowheads) and the VHP from P8-P10 (Fig. 4D-F). Connections between VHP vessels and the TVL were nevertheless amenable for investigation (Fig. 4Z, magenta arrows - TVL rupture, yellow arrow - VHP-TVL connection point. Ritter et al demonstrated completion of hyaloid regression by P46^6^, however we observed complete regression (aside from isolated vessels segments) at P28 (Fig. 4W). In contrast, Darche et al showed remaining vessels still connected to the optic nerve at P60 in albino eyes^5^. This discrepancy might be explained by difference in mouse strains used. The last stage we observed hyaloid vessels connected to both the optic nerve and iris was P20 (Fig. 4V, Suppl. Fig. 4A, B).

**Figure 4.**
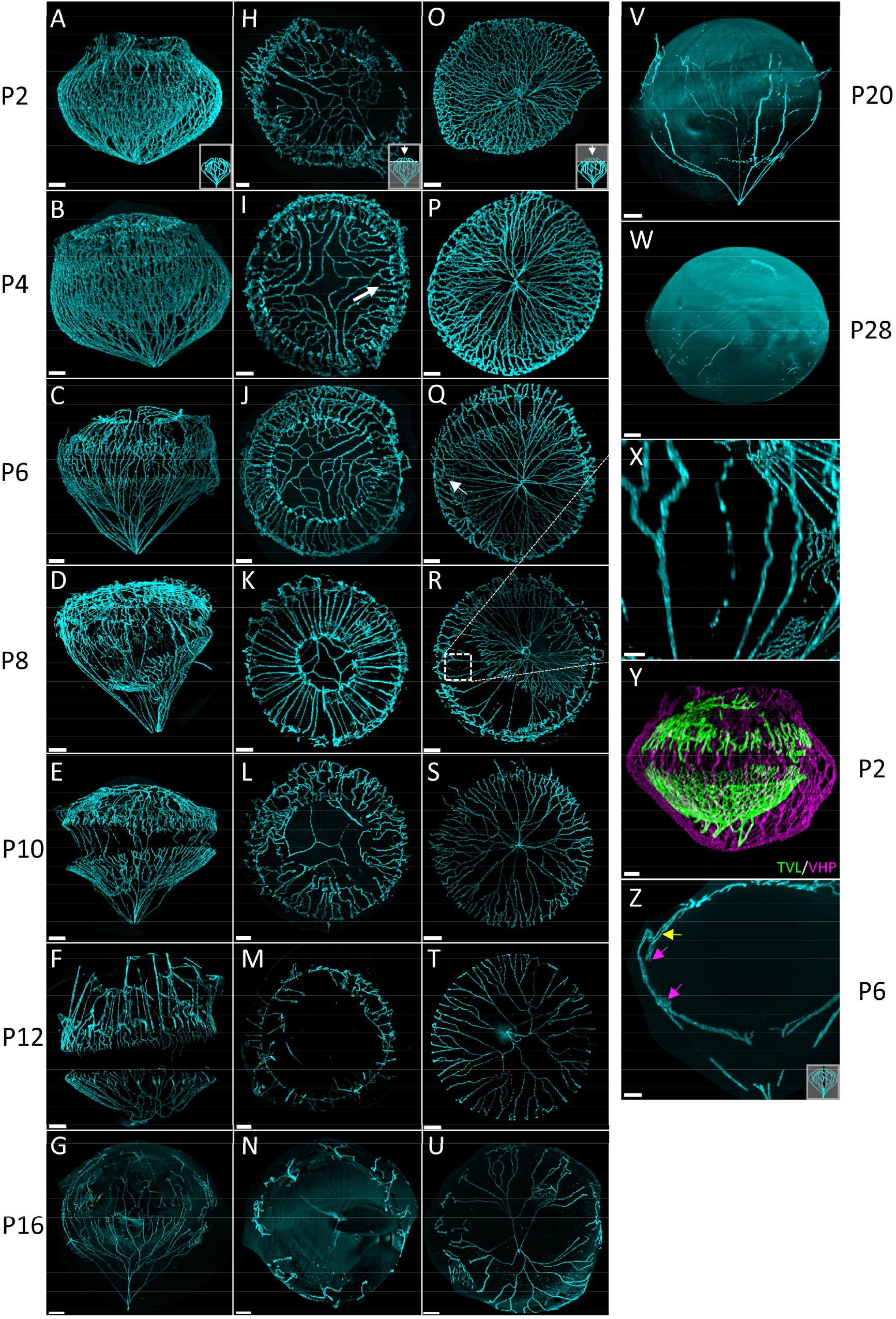
Regression of the postnatal hyaloid vasculature. (**A-G**, **V, W**) Side view of the entire hyaloid system during its regression from P2 up to P28. (**H-N**) Anterior view of the regressing PM and anterior TVL (lower hyaloid not displayed). (**O-U**) Anterior view of the lower hyaloid system (PM and anterior TVL is hidden). (**I**) Arrow indicates first signs of vessel fragments in the PM at P4. (**X**) Higher magnification of disintegrating vessels. (**Y**) In silico dissected VHP (magenta) and TVL (green) at P2. (**Z**) Cross section of P6 hyaloids, the VHP connects to the vessels of the TVL on the top of the lens. The TVL is ruptured with the lens (magenta arrows). Labelling: CD31. Scale bars: A-C, I, J, N-Q, T = 150 μm, D-G, K-M, R, S, U-W = 200 μm, H, X-Z = 100 μm.

### Remodelling of the developing iris vasculature visualised in 3D

The vasculature of the iris is exposed to continuous stretching and compaction as a result of light adaptation. Although the iris vasculature has been well characterised in adult conditions, little is known of its postnatal development and early morphogenesis. At birth the iris receives blood flow from the VHP and TVL (Fig. 1L), as well as from the long posterior ciliary arteries that also supply the ciliary body vasculature (Fig. 1M)^27^. From the iris, blood travels to the choriocapillaris, that drain into the vorticose veins (Suppl. Fig. 1C, F). Regression of the hyaloids would impose dramatic changes in perfusion of the iris vasculature in turn affecting its remodelling. To study the iris vasculature around the period of hyaloid regression we analysed LSFM-acquired data from various time points. At P4, the iris vasculature connected to both the hyaloid and PM vessels, thereby often forming intersections of three vessels (arrow, Fig. 5A, angled view Suppl. Fig. 5A, Suppl. Movie #3). However, single connections to either PM or hyaloid vasculatures were also evident. At P8, the iris vasculature was primarily connected to hyaloid vessels, although some intersections with the PM remained. Interestingly, some iris vessels extended over the pupil, before connecting to the remaining PM-vasculature (Fig. 5B arrows, angled view Suppl. Fig. 5B). Subsequently, at P12, loops of vessels were observed at the edge of the iris, possibly by fusion of iris intrinsic vessels (arrows, Fig. 5C). Following the hyaloid-iris disconnection, the iris vasculature further remodelled as indicated by differences between the perinatal and adult iris vasculatures (Fig. 5D).

**Figure 5.**
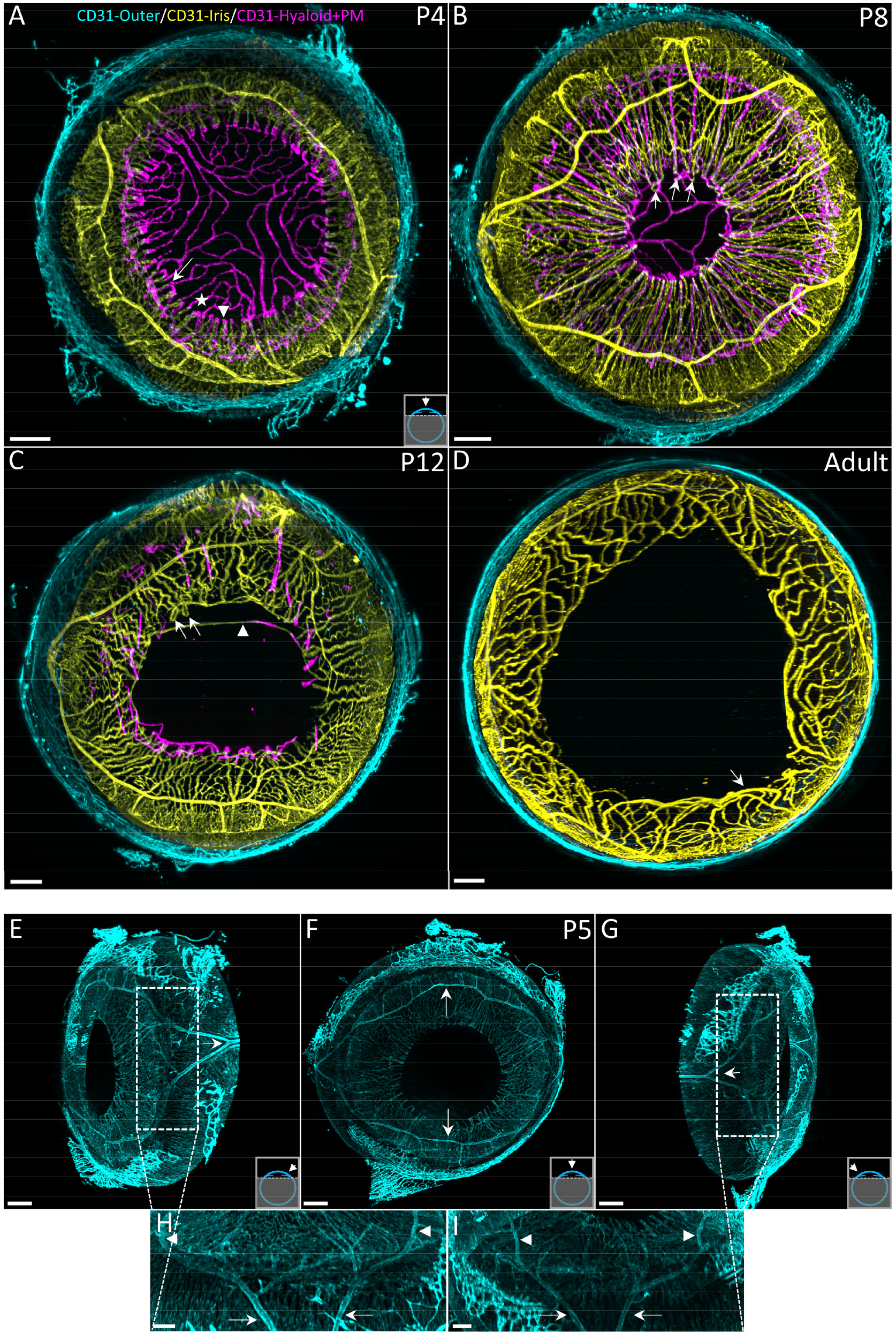
Postnatal development of the iris vasculature. (**A-D**) Iris vasculature (yellow), anterior hyaloid and PM vasculature (magenta), uveal and limbus vasculature (cyan), all CD31 labelled. (**A**) Anterior view of a P4 eye with dilated pupil shows extensive PM network. Most hyaloid-iris vessel connections are intersected by PM vessels (arrows), though some are uninterrupted (arrowhead). PM-to-iris vessels also exist (star). (**B**) A P8 eye with substantially regressed PM network. Some remaining vessels may hide posterior to the constricted iris. Most hyaloid-to-iris vessels are still present, of which some extend over the pupil (arrows). (**C**) In the P12 eye the PM network is gone, except for one remaining vessel (arrowhead). Arrows indicate atypical looping iris vessels. (**D**) An adult eye in which the PM and hyaloids have completely regressed. Vessels pointing towards the pupil are arching away instead of going over the pupil (arrow). (**E-G**) P5 eye showing the long posterior ciliary arteries, bifurcating at the limbus area (arrows, **E, G**) to supply the iris. Arrows indicate anastomosing iris arteries (**F**). Side views of the ciliary-to-iris arteries. (**H, I**) close-up of ciliary artery(arrows)-to-iris artery(arrowheads) transition. Labelling: CD31. Scale bars: A-D = 200 μm, E-G = 150 μm, H, I = 100 μm.

The bifurcated ciliary arteries were observed to enter the iris at both sides of the eye (Fig. 5E&G, enlarged in H, I). In several samples, across developmental stages, the ciliary arteries anastomosed in the middle of the iris (arrows Fig 5F, Suppl. Movie #4), implying arterial blood flow in opposing direction (P4 Fig. 5A, P8 Fig. 5B, P16 Suppl. Fig. 5C-F). These anastomosing arterial connections appeared transient as they could not be observed in adults. Although the iris vasculature is considered to drain into the choriocapillaris, the vascular connections have to our knowledge not been illustrated in 3D. Using optical clipping planes in Imaris, the intricate connections between the iris vasculature and the choriocapillaris could be visualised and followed towards the vorticose vein (Suppl. Fig. 5H-J, Suppl. Movie #5). Although a full characterisation of these dynamic events can only be achieved through longitudinal intravital imaging, we demonstrate here that details of vascular remodelling and blood flow in the interface of the hyaloid, PM and the iris vasculature can be resolved with LSFM of intact mouse eyes.

### Analysis of laser-induced choroidal neovascularisation

As demonstrated above, LSFM of cleared eyes allowed for in depth 3D analysis of all vasculatures during postnatal development, and in adulthood in a physiological context. As proof of concept for utilisation of this technique in studies of eye pathology, we turned to the biology of neovascular age-related macular degeneration (nAMD). In nAMD, angiogenic choroid vessels penetrate the Bruch’s membrane and the RPE, leading to photoreceptor atrophy and vision loss. This critical stage of neovascularization can be mimicked by laser-inflicted wounding of Bruch’s membrane in the mouse ^28^. Post-mortem microscopy-based analysis of such tissue has so far been limited to sections of eyes or whole-mount preparations of individually dissected choroids or retinas^29^. Here, we combined laser-induced choroidal neovascularization (CNV) with the LSFM workflow, for evaluation of analytical potential.

Mice exposed to laser-induced wounding of the Bruch’s membrane were followed live by stereoscopic observations (Fig. 6A), optical coherence tomography (OCT) (Fig. 6B, arrow), as well as by fluorescence angiography (FA), exposing potential leakage within lesions (Fig. 6C, star), until sacrifice at day 7. Then, enucleated eyes were run through the LSFM workflow. 3D analysis of CD31-stained structures permitted matching of individual retina and choroid lesion-sites to those previously imaged *in vivo* by OCT/FA (Fig. 6A-F). Positioning of the lesion perpendicular to the camera in LSFM, provided acquisition of subcellular details throughout the entire lesion (Fig. 6G-I, Suppl. Movie #6), at a resolution beyond what can be achieved by OCT (Suppl. Movie #7, 8).

**Figure 6.**
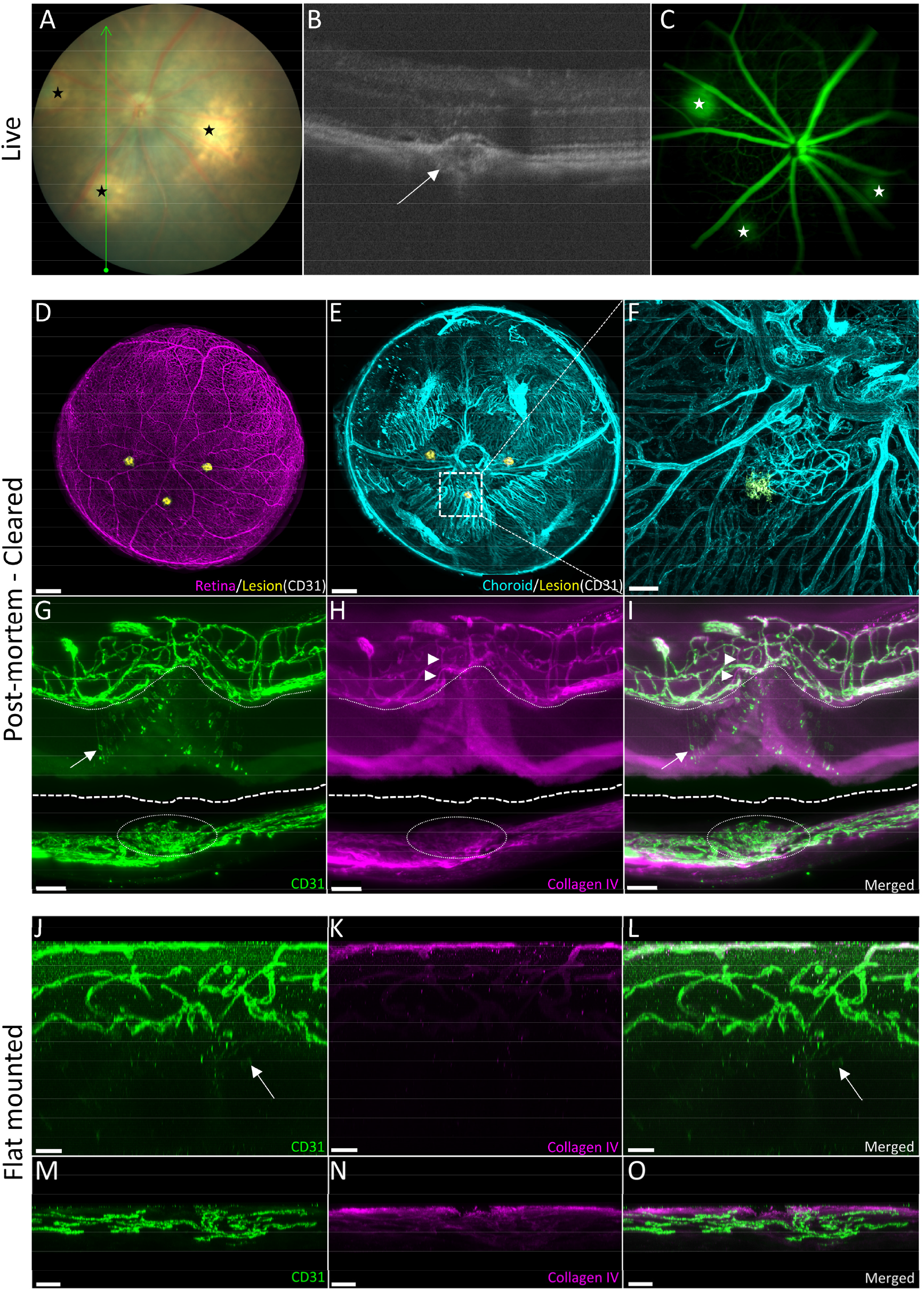
The LSFM workflow provides new perspectives on laser induced choroidal neovascularisation (CNV). (**A**) Stereoscopic view of the retina/posterior eye in vivo at 7 days (D7) post CNV induction. Stars indicate lesions. Green line indicates the optical plane visualised by Optical Coherence Tomography (OCT) in **B**. (**C**) Stereoscopic view of fluorescence angiography, displaying leaky lesions (star), on D7 post CNV induction. (**D, E**) The same CNV-induced eye post-mortem, following immunolabelling for CD31 and αSMA, tissue clearing and LSFM. The retina (**D**) and choroid (**E**) were in silico dissected, as well as the lesions indicated in yellow. (**F**) High resolution LSFM acquisition of the square in **E** (the muscle vasculature was omitted for clarity). (**G-I**) Cross section (100 μm) of a D7 CNV lesion, immunolabelled with CD31 (**G**) and Collagen IV (**H**, merged **I**). Retina and choroid are separated by in silico dissection (dashed horizontal line), for optimization of intensity in sub layers. Note CD31+ cells and empty collagen sleeves in the retina (arrowheads). The choroid vascular lesion is highlighted by the dotted ellipse. The retinal vasculature appears pushed up (upper dotted line). (**J-L**) Confocal microscopy-acquired cross section (100 μm) of lesion site of the same eye, following relabelling and flat-mounting, stained for CD31 (**J**) and Collagen IV (**K**, merged in **L**). CD31+ cells in the ONL can be observed (arrows J&L), though much less clear than in G&I. (**M-O**) Confocal microscopy-acquired cross section (100 μm) of lesion site following relabelling and flat-mounting of the choroid and RPE. CD31 (**M**) and Collagen IV (**N**, merged in **O**). Scale bars. D, E = 300 μm, F = 100 μm, G-I = 50 μm, J-O = 15 μm.

Microscopy of flat-mounted tissue is routinely applied for high-resolution analysis of laser-induced CNV in mice. To compare LSFM and confocal-assisted analysis of CNV lesions, we first studied CD31 and collagen IV immuno-stained cleared whole-eyes with LSFM (Fig. 6G-I) and then physically dissected the retina and choroid for re-staining, flat-mounting and confocal microscopy (Fig. 6J-O, Suppl. Fig. 6). Confocal microscopy enabled superior resolution to LSFM in the anterior view (x-y) of the lesion (Suppl. Fig. 6 vs Fig. 6F), but inferior resolution in the cross-sectional view (Fig. 6G-I vs Fig. J-O). Flat-mounting also disturbed the 3D architecture of the tissue (Fig. G-I vs Fig. J-L). In addition, recording of structures within the deeper layers was drastically worse with laser scanning confocal microscopy compared to LSFM, in particular at lower wavelengths (Fig. 6H vs Fig. 6K, N). The LSFM workflow also exposed previously undescribed CD31+ positive cells with distinct soma and protrusions within the outer nuclear layer (Fig. 6G&I & Suppl. Fig 6G-I), which will merit further studies herein. Here we show that LSFM of CNV lesions in cleared intact eyes improves on many technical obstacles that currently limit the understanding of CNV progression.

## Discussion

Immuno-labelling, depigmentation, optical clearing, and LSFM of intact mouse eyes provide new opportunities for visualisation and analysis of eye anatomy in development and disease. The eye is particularly well-suited for this approach as it is compatible with in silico dissection into volumes of interest due to its compartmentalised anatomy. Thus, specific data can be acquired for individual vascular beds even with single antibody labelling thereby exposing potential tissue crosstalk. As presented herein, all vasculatures of the mouse eye can be assessed in high detail, without physical dissection. For follow up analysis at even higher resolution, we provide a workflow for consecutive physical dissection and flat-mounting of compartments. The workflow is applicable to any developmental stage or mouse strain and is well suited for analysis of disease models as demonstrated by novel descriptions of the laser-induced CNV model. Hence, optical clearing and LSFM of intact eyes overcomes technical challenges and provide a whole-organ context in the study of development and complex eye disease.

In recent years, the application of tissue clearing and LSFM for the study of ocular vasculature has seen major advances. Several groups have described clearing of ocular tissue to facilitate imaging, with Singh et al^12^ being the first to apply LSFM on cleared dissected retinas (rat) and quantification in 3D. Later, Prasht et al used a similar approach of resection of the sclera and RPE and demonstrated the benefits of tissue clearing over traditional flat-mounting in terms of preserved vessel dimensions^13^. The first clearing of whole eyes was performed by Henning and colleagues in the study of healthy adult retinal vasculature and main choroid arteries, also demonstrating the use of the latter for anatomical orientation^9^. Vigouroux et al applied whole-eye clearing on transgenic mouse models in the study of eye development, showing its power in such a context, though they did not study the vasculature^15^. Yang et al were first to display several of the ocular surface vasculatures through the CLARITY protocol^10^. Gurdita et al^11^ used LSFM on CUBIC-cleared eyes of several transgenic and disease models, following the proof of principle by Ye et al^30^. Finally, Darche and colleagues described the regression of the hyaloid vasculature using LSFM on intact albino eyes^5^. Extending on previous work we present the first compilation of all eye vasculatures of postnatal and adult mice. Detailed visualisation of the connectivity of the various vasculatures at different developmental stages exposes their dynamic nature. This includes pruning and disconnections of the hyaloids, a process expected to have direct impact on flow patterns and remodelling of the remaining vasculatures of the eye. These features highlight the relevance of extending beyond traditional analysis of individual sub-vasculatures – such as the retina – to include other vasculatures with potential dramatic impact on eye physiology.

The clearing protocol described here is a combination and adaption of the previously published DEEPClear, EyeCi and eyeDISCO protocols^9,14,15^ and applies RI-matching (clearing) and immunostaining in accordance with the rapid and easy iDISCO+ approach^16^. The resultant protocol has a sample processing time of 10 days for full visualisation of all vasculatures of the eye, and does not require antibody injection or enzymatic digestion, as applied in similarly fast protocols ^9,11,15^. Selection of the best suited workflow for an individual sample relies on several factors, such as availability of imaging equipment, antigen/antibody compatibility, the use and need of endogenous fluorescent reporter preservation, pigmentation, tissue composition, sample size, timeline et cetera. The protocol presented herein does not preserve endogenous fluorescence, although application of antibodies to specific endogenous reporters is possible. Preservation of tissue morphology is important and while our workflow presents with the caveat of distortion of the lens thereby affecting the VHP and TVL, most vasculatures can be studies in great detail in their entirety. However, other protocols have shown minimal effects on tissue morphology and compose valuable alternatives for certain experimental settings ^5,11^. Further development of protocols to retain eye shape, fast bleaching and immunolabelling and good antigen preservation are warranted. Irrespective of what method has been used to facilitate data collection by LSFM, we and others demonstrate that in silico dissection provides great opportunities for analysis and visualisation of individual vascular beds of the eye in a complete organ context^5,11^.

Whole-eye imaging promoted identification of temporal-nasal and dorsal-ventral anatomy in turn facilitating characterisation of anatomy-matched asymmetrical radial- and deep layer expansion of the postnatal retinal vasculature. Why dorsal progression is slowest, and ventral fastest is still unexplained, but the underlying cause may contain critical information on mechanisms of vascular development and patterning. This further exemplifies the added value of analysing the eye in its entirety.

Due to its inherent fragility, the transient hyaloid system has been difficult to study. Physical dissection, histological sectioning and electron microscopy have provided fundamental information on hyaloid biology but have generated limited knowledge on the interplay with other vasculatures in remodelling. Similar to Darche et al^5^, we present a workflow by which successive regression of the hyaloid vasculature, in relation to the other vasculature, can be followed thereby adding to the relevance of the strategy in studies of the biology of human eye disease.

Simultaneous to the process of hyaloid regression, the iris vasculature undergoes extensive remodelling. Despite the iris vasculature being accessible for microscopy, its postnatal development has been poorly described. In silico dissection of whole eye vasculature data into iris, hyaloid and PM compartments allowed for simultaneous assessments of their connections, as well as their individual anatomy. The analysis demonstrated extensive remodelling of the iris vasculature in conjunction with, and likely as a response to, hyaloid regression. In addition, the whole-eye perspective enabled analysis of the iris blood supply and drainage, revealing the existence of anastomosing arteries on the iris surface in some postnatal eyes. Although analysis of fixed samples provides great detail on differences over time, as presented here, only longitudinal intravital imaging can fully resolve the dynamic properties of individual vessels and their flow^6,31,32^.

Finally, our application of laser-induced CNV as a mouse model for nAMD in combination with LSFM allowed for whole-lesion imaging thereby promoting novel observations of the relations of the retinal and choroidal vasculatures. These observations could not be made on physically separated or flat-mounted samples. Also, in the cross-sectional plane, LSFM provided better resolution than standard confocal microscopy. These features highlight the potential of LSFM in the study of models of complex eye diseases.

For numerous reasons, LSFM data of cleared whole organ samples is difficult to quantify^33^. Light from the laser as well as the fluorophores lose brightness and may be distorted as it travels through tissue, resulting in regions of bright and sharp along with dim and vague signal in the same sample. Intensity and contrast-based measurements such as volumes, cell counts, surface areas, etc. can therefore be unbalanced. The asymmetry of the eye adds further complexity to this problem. To date implementation of manual supervision and adjustments of signal intensities are required to utilize samples optimally.

In conclusion imaging intact eyes followed by in silico dissection and detailed analysis maximises data collection from a single mouse. Here we demonstrate the power of LSFM on cleared whole eyes by providing a compilation of all vasculatures of the developing and adult mouse eye. Likely application of LSFM on intact eyes will provide new perspectives in developmental as well as pathological ophthalmology, enabling advancements in understanding towards improved therapies.

## Acknowledgements

This study was supported by the Swedish Research Council (2021-01210 to L.J.; 2021-03108 to P.U.), the Swedish Cancer Society (190487 Pj to L.J., CAN 19 0544 Pj and 19 0545 Us to P.U.), the Swedish Childhood Cancer Foundation (PR2020-0124 to P.U.), the Swedish Brain Foundation (FO190101 to L.J.; FO2021-0230 to P.U.), the Cancer Research Foundations of Radiumhemmet (grant 201362 to P.U.), The Swedish Heart and Lung Foundation (20190531 to L.J.), Erik Stenbäck Foundation (to L.J.), The Swedish Eye Foundation (Ögonfonden, to L.J. and H.A.), a European Union’s Horizon 2020 research and innovation programme under the Marie Skłodowska-Curie grant agreement (No 814316 to L.J.) and Karolinska Institutet. The light-sheet microscopy infrastructure used in this research received grants from the Strategic Research Area in Neuroscience (StratNeuro) and the Strategic Research Area in Stem Cells and Regenerative Medicine (StratRegen), supported by the Swedish government. We thank the BIC imaging core-facility for the use of their light-sheet microscope and image analysis workstation.

## Author contributions

L. Jakobsson, L. Krimpenfort, H. André and P. Uhlén designed the research study; L. Krimpenfort, M. Garcia-Collado, T. van Leeuwen, F. Locri, AL. Luik, A. Queiro Palou, S. Kanatani conducted experiments and acquired data; L. Jakobsson, L. Krimpenfort, T. van Leeuwen, F. Locri, and M. Garcia-Collado analysed data. All authors contributed to the writing of the manuscript.

