## Supplemental info and figs for "Anatomy of the complete mouse eye vasculature in development and pathology explored by light-sheet fluorescence microscopy"

### Supplemental information

Contents:
- Supplemental figures with legends (P2-7)
- Supplemental movie legends (P8)
- Supplemental materials and methods (P9-11)
- Tutorial for Imaris-based analysis of whole mouse eye vasculature using in silico dissection (P12-20)

#### Supplemental figures with legends

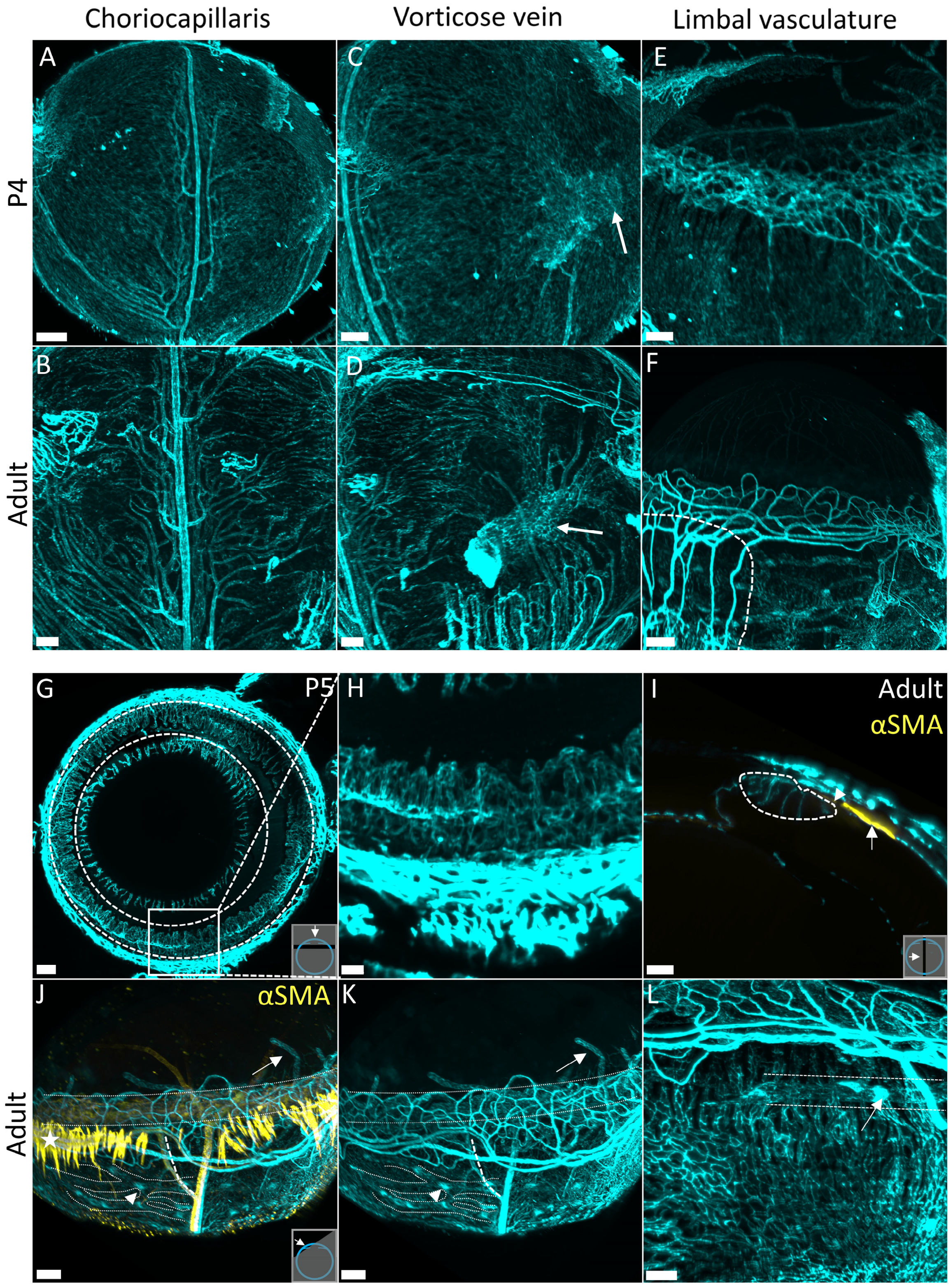


**Supplemental Figure 1. The choroid artery, choriocapillaris, vorticose vein, limbal vessels, Schlemm's canal and lymphatic vasculature.** (**A, B**) Lateral choroid artery, vessels and choriocapillaris at P4 and adult stage. (**C, D**) vorticose vein at P4 and adult stage. Due to enucleation of the eye, the part of the vessel outside the sclera is removed. (**E, F**) limbus vasculature at P4 and adult stage. The limbus vasculature is connected to the eye-muscle vasculature (dotted line, **F**). (**G**) frontal cross-section (20 μm) of the ciliary body vasculature at P5 (between dotted circles). (**H**) enlarged view of ciliary body vasculature from box in **H**. (**I**) Horizontal cross section (3 µm) of ciliary body vasculature (dotted oval) at adult stage. αSMA staining indicates the ciliary muscle (arrow) and Schlemm's canal (arrowhead). (**J, K**) Side view of the anterior eye immunolabeled for αSMA (**J**, yellow) and CD31 (**J, K**, cyan). Schlemm's canal is highlighted by dotted lines, star indicates ciliary muscle, arrow indicates blind-ended lymphatic vessels, dashed line indicates branch of ciliary artery feeding the limbus vasculature (**J, K**), arrowhead points to lymphatic valves of lymphatic collecting vessels highlighted by dotted lines. (**L**) Closer view of a lymphatic vessel, potentially draining into the choriocapillaris. Lymphatic valve indicated by arrow. Labelling: CD31, unless otherwise specified. Scale bars, A, B, D, J-L = 200 µm; C, E = 150 µm; F = 300 µm; G, I = 100 µm; H = 50 µm.

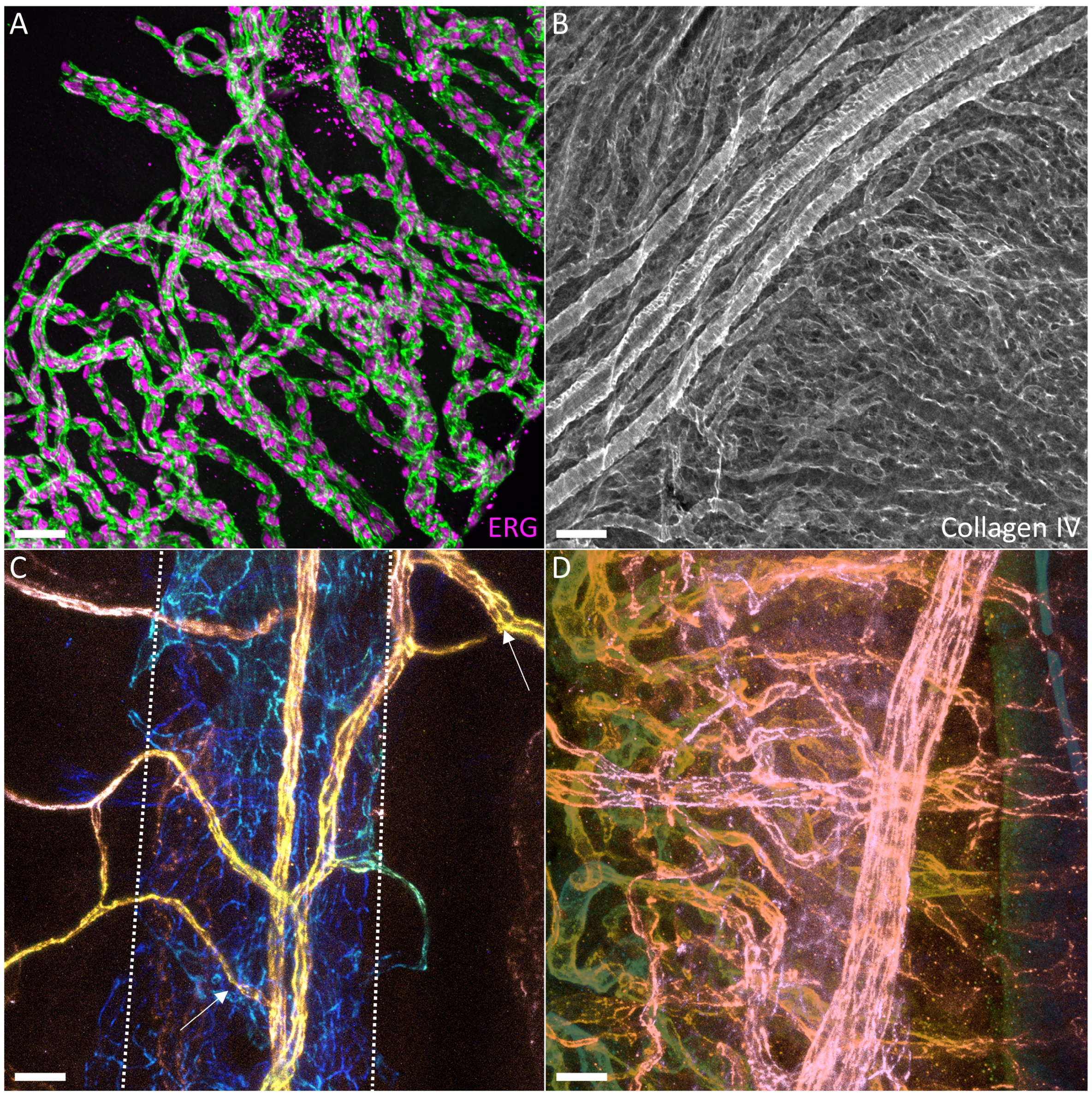


**Supplemental Figure 2. High resolution confocal acquisitions of previously cleared tissues.** (**A**) Hyaloid vessels immunolabeled for CD31 (green) and ERG (endothelial cell nuclei, magenta). (**B**) Choroid vasculature immunolabeled for collagen IV. (**C**) Schlemm's canal (dotted lines) and limbus vasculature (arrows), depth colour coded. (**D**), Ciliary body vasculature, depth colour coded, (**C&D**) derived from same image stack, separated for clarity. Labelling: CD31, unless otherwise specified. Scale bars = 30 µm.

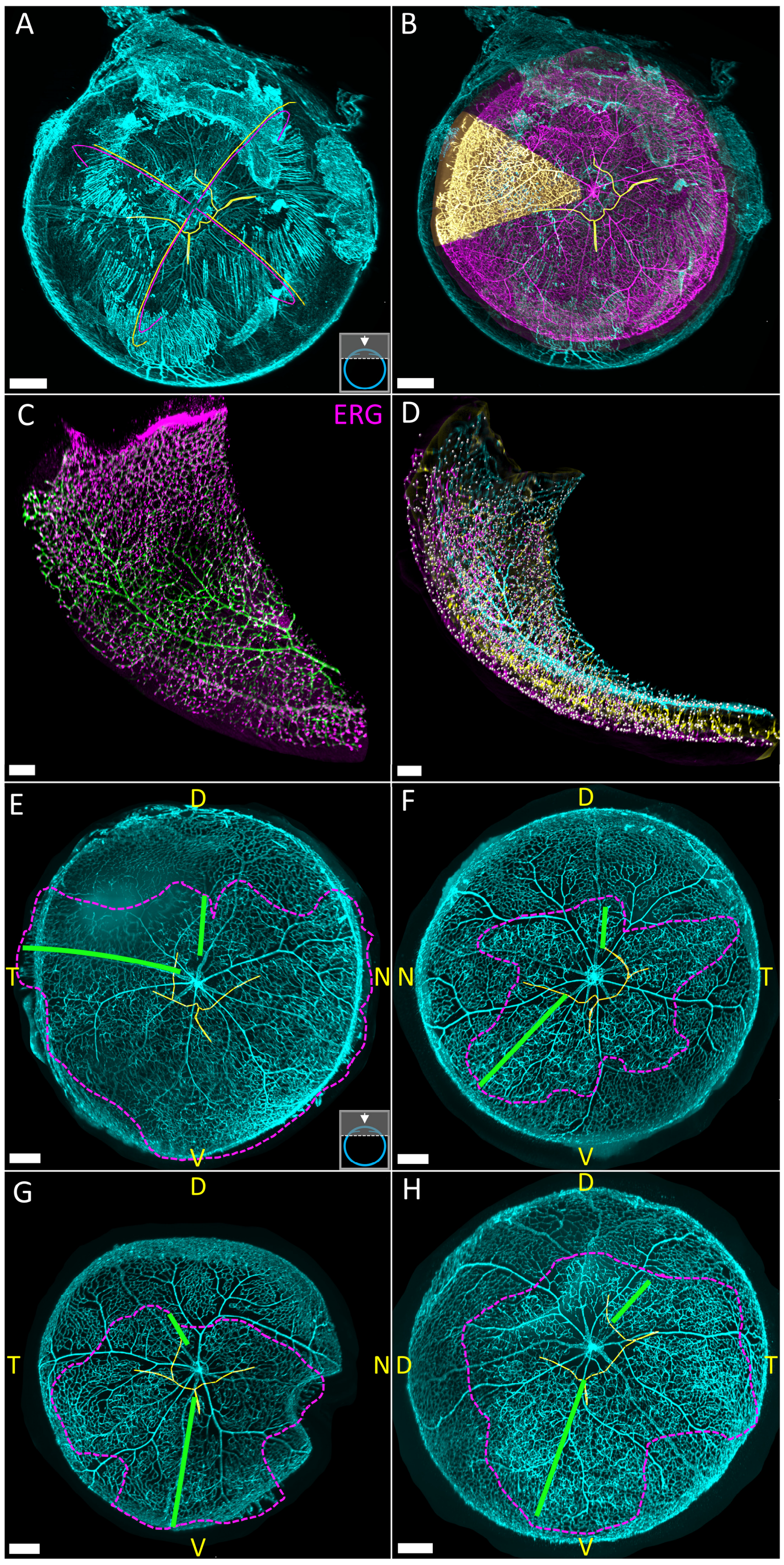


**Supplemental Figure 3. Analysis of asymmetrical development of the deep retinal vascular plexus with anatomical correlation.** (**A**) Measurement of superficial and deep plexus establishment. Magenta and yellow lines indicate superficial and deep plexus measuring lines respectively. (**B**) Isolation of part of the retina (yellow slice) for further analysis, care was taken to include one artery and one vein. (**C**) Isolated part of the retina, immunolabelled for CD31 (green) and ERG (magenta). (**D**) Spots (white) analysis for comparison of cell number between the superficial (cyan), intermediate (yellow) and deep plexus (magenta). (**E-H**) P8 retinas used for measurement of shortest and longest (green lines) deep plexus establishment. Orientation of choroid arteries and rough outline of deep plexus establishment indicated in yellow and (dashed) purple lines. V = ventral, T = temporal, N = nasal, D = dorsal. Labelling: CD31, unless otherwise specified. Scale bars, A,B = 300 μm, C,D = 100 μm, E-H = 200 μm.

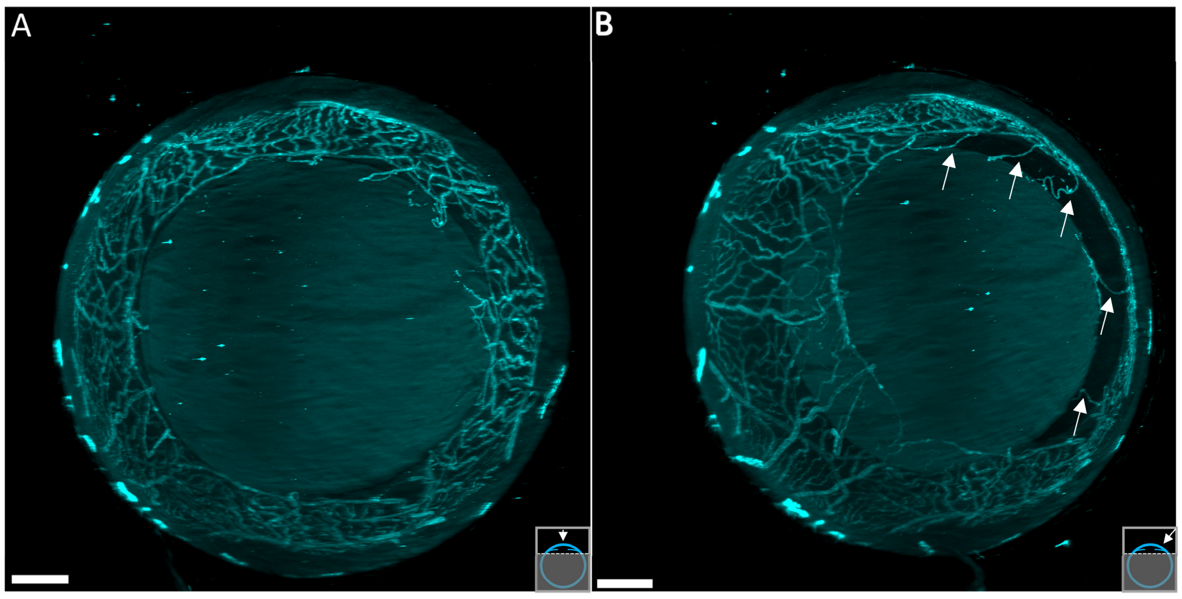


**Supplemental Figure 4. Hyaloids at P20 are connected to the iris vasculature in some eyes.** (**A**) Anterior view of the iris at P20, immunolabelled for CD31. (**B**) Slanted view of the same iris, indicating the remaining hyaloids (arrows) are still connected to the iris vasculature. Scale bars = 300 μm.

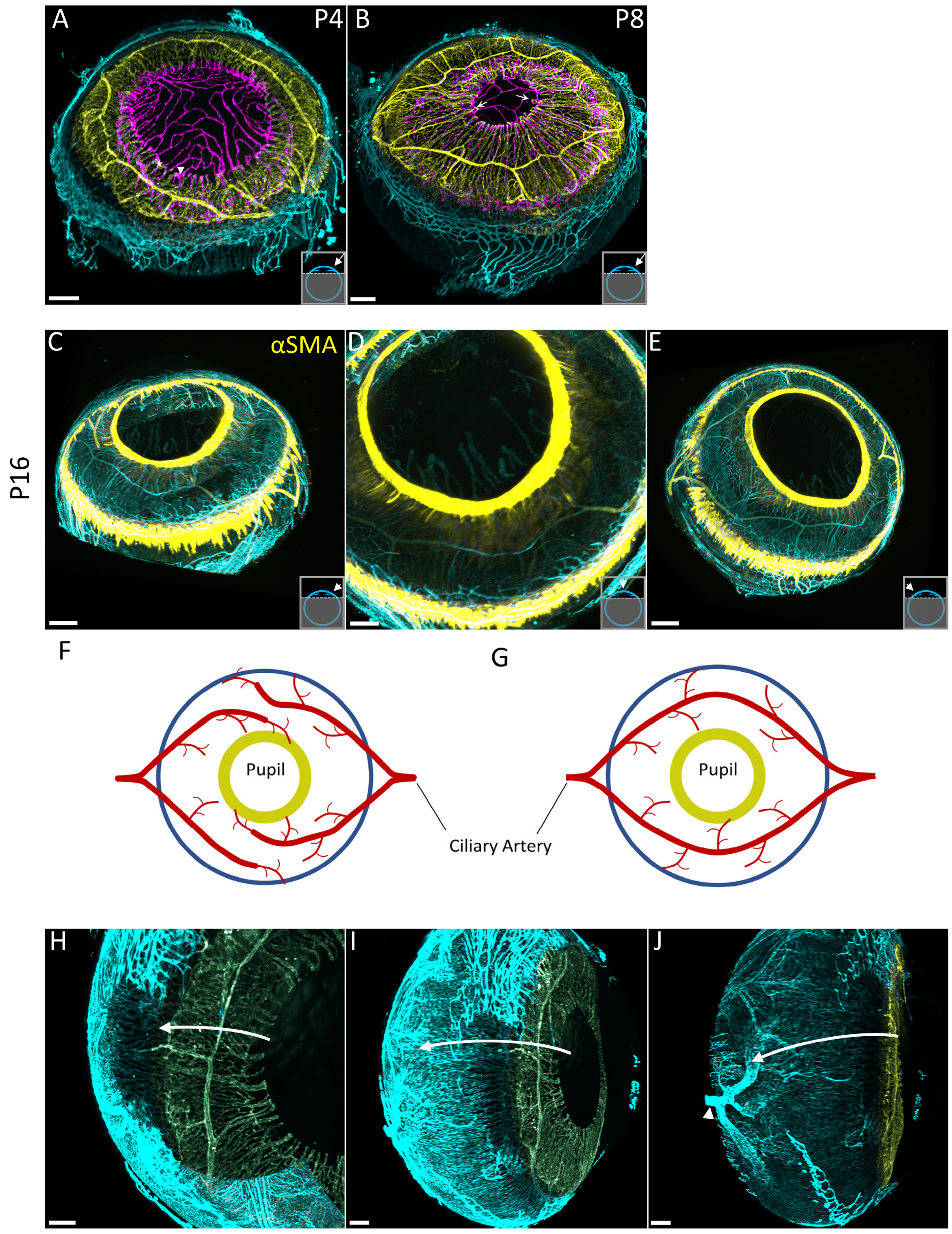


**Supplemental Figure 5. Vascular connections and blood flow of the iris.** (**A&B**) Side view of anterior eyes shown in figure 5 A&B. (**C-E**) The anterior of a P16 eye immunolabelled for CD31 (cyan) and αSMA (yellow), showing anastomosed ciliary-to-iris arteries. (**F**) Schematic of commonly observed iris blood supply, with the ciliary arteries supplying the arteries in the iris, which diverge before connecting. (**G**) Schematic demonstrating occasional observations of anastomotic ciliary artery connections in the developmental iris. (**H-J**) Drainage of the iris vasculature happens over its entire circumference into the choriocapillaris (**H&I**), which subsequently drain into the vorticose vein (**J**, arrowhead). Direction of blood flow indicated by arrows, inferred from anatomy (**H-J**). Labelling: CD31, unless otherwise specified. Scale bars, A,B,D = 200 μm, C,E = 300 μm, H-J = 100 μm.

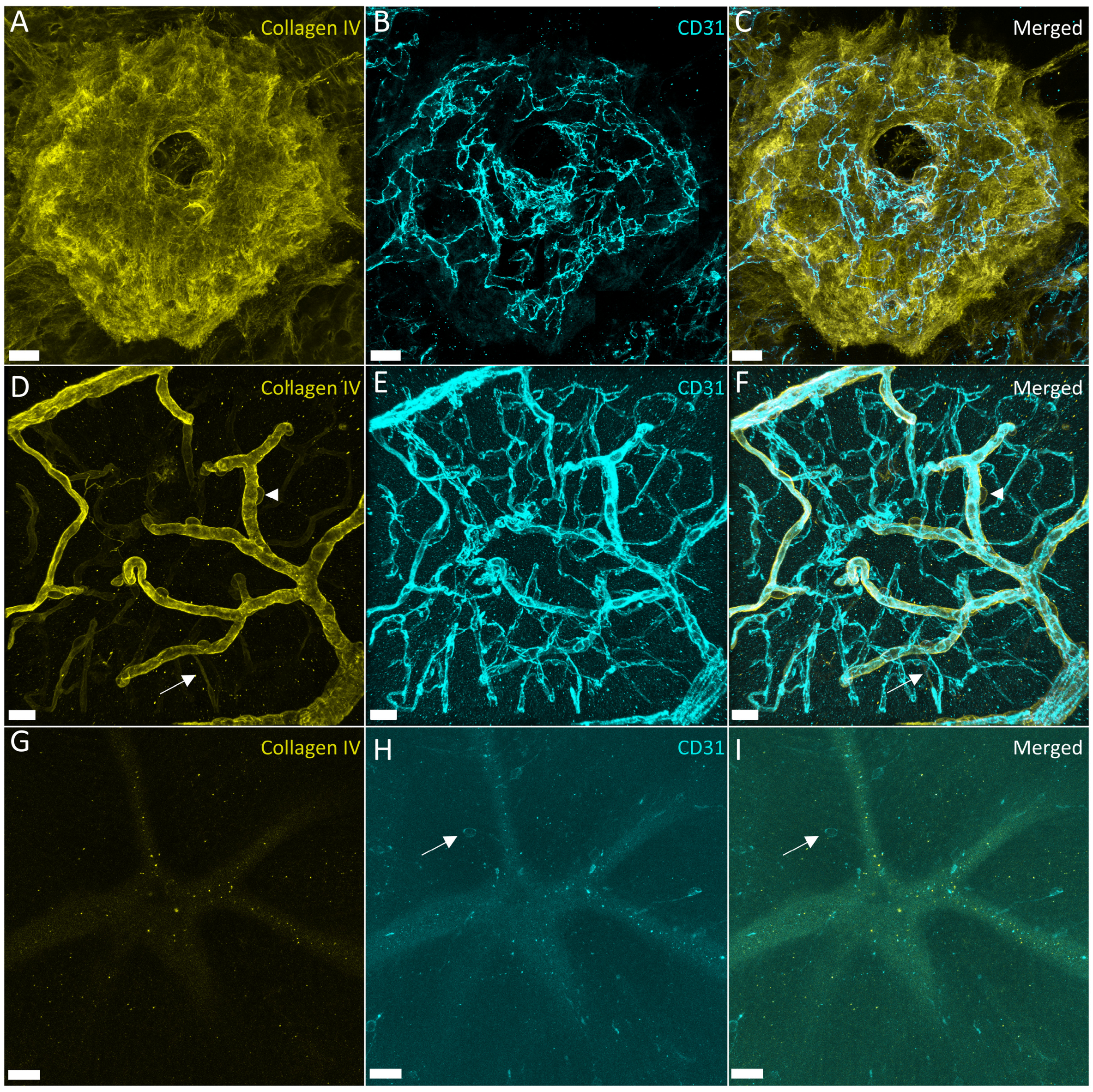


**Supplemental Figure 6. Confocal acquisitions of CNV lesions of RPE and retina flat mount, 7 days post laser induction.** (**A-C**) Cross section (20 μm) of CNV lesion site on RPE flat mount immunolabelled for collagen IV(**A**) and CD31 (**B**, merged in **C**). (**D-F**) CNV lesion site within the flat-mounted retina, immunolabelled for collagen IV (**D**) and CD31 (**E**, merged in **F**). Collagen encasing mural cells (arrowheads) and empty collagen sleeves (arrow) can be observed. (**G-I**) CNV lesion site on retina flat mount, omitting the vascularized layers, immunolabelled for collagen IV (**G**) and CD31 (**H**, merged in **I**). CD31+ cells can be observed (arrows). Scale bars, A-F = 20 μm, G-I = 30 μm.

#### Supplemental movie legends

**Supplementary Movie #1. In silico dissection of the eye reveals the different sub compartments by differential pseudo-coloring.** 0:00-0:04, 2D-stack of whole-eye light-sheet acquisition, immunelabelled for CD31. 0:05-0:10, 3D rendering of the whole-eye. 0:11-0:13, isolation of the inside eye using the grey volume, revealing the retina and hyaloid vasculature. 0:14-0:24, isolation of the hyaloid vasculature using another volume, enabling visualisation of the retina, hyaloid and outer vasculature in different colours (yellow, magenta, cyan). 0:25-0:30, details in the dense choroid can be observed. 0:32-0:35, a clipping plane obscures the posterior eye, revealing the iris vasculature. 0:35-0:39, in silico dissected iris is made yellow, and magenta hyaloid and PM vasculature is brought back. 0:40-0:45, GLUT-1 staining in cyan shows the structure of the iris. 0:46-0:50, overview of the outer, retinal and hyaloid vasculatures (cyan, yellow, magenta).

**Supplementary Movie #2. The hyaloid and retinal vasculature.** An Imaris clipping plane enables progressive visualisation of the retinal and hyaloid vasculature from the optic nerve to the anterior lens. The retina and hyaloid vasculature derive their blood supply from the same artery, but the retina drains through the optic nerve whereas the hyaloids drain via the iris at the anterior lens.

**Supplementary Movie#3. Hyaloid-to-iris connections.** 3D visualisation of the outer, iris and hyaloid vasculatures. Close-up of the iris-hyaloid interface shows the connections between hyaloid, iris and PM vessels.

**Supplementary Movie#4. Clipping planes help reveal anastomosed iris arterial blood supply.** 0:00-0:05, outer and iris vasculature, iris arteries run laterally on the top and bottom of the iris, but the origin of the arteries is hidden by the limbus vasculature. 0:05-0:10, use of clipping planes omits the limbus vasculature, revealing the long posterior ciliary arteries that supply the anastomosed iris arteries from both sides.

**Supplementary Movie#5. Drainage of the hyaloids, via the iris and choriocapillaris to the vorticose vein.** 0:00-0:05, hyaloid-to-iris drainage. 0:05-0:11, drainage of the iris is hidden again by the limbus. 0:12-0:20, omission of limbus vasculature by use of clipping planes enables tracing of anatomy inferred blood flow routes from the iris to the vorticose vein.

**Supplementary Movie#6. CNV lesion imaged by LSFM.** High-resolution acquisition enables detailed visualisation of the entire CNV lesion in a cross-sectional manner. Labelling: CD31=green, Collagen-IV = magenta. Choroid and retina are in silico dissected to allow optimal contrast settings for both.

**Supplementary Movie#7. 3D-visualisation of OCT acquired lesion.** 0:00-0:08, focus on choroidal side of lesion. 0:08-0:11, choroid and retina viewed together. 0:11-0:17, focus on retinal side of lesion. 0:18-0:20, anterior view of choroid lesion.

**Supplementary Movie#8. 3D-visualisation of LSFM acquired lesion.** 0:00-0:06, focus on choroidal side of lesion, lesion highlighted in yellow at 0:03. 0:07-0:14, focus on retinal side of lesion. 0:15-020, posterior view of choroidal lesion.

#### Supplementary material and methods

| Reagent | Company | Product number |
|---|---|---|
| **DBE** | Aldrich | 108014 |
| **DCM** | Sigma | 270997 |
| **Donkey Serum** | Jackson Immunoresearch | 017-000-121 |
| **Triton X-100** | Sigma | T-8787 |
| **THEED** | Aldrich | 87600 |
| **Acetone** | Sigma-Aldrich | 179124 |
| **Methanol** | Sigma-Aldrich | 34860 |
| **Gelatin from porcine skin** | Sigma | G1890 |
| **Glycine** | Merck | 1.04201.100 |
| **Heparin** | Sigma | H-3149 |
| **Tween-20** | Sigma Aldrich | 102423676 |
| **PFA** | Sigma Aldrich | P6148 |
| **Urea** | Sigma | U5378 |
| **DMSO** | Thermo Scientific | 295520010 |
| **30% $H_2O_2$** | Sigma | H1009 |
| **PBS10X** | Medicago AB | 12-9423-5 |
| **BSA** | Saveen Werner AB | B2000-100 |
| **Prolong Gold** | ThermoFisher Scientific | P36930 |
| **Isoflurane** | Baxter | VDG9623C |
| **Tropicamide (0.5%)** | Alcon | 0998-0355-15 |
| **Saline (9mg/mL NaCl)** | Fresenius Kabi | 210352 |
| **Viscotears** | APL | 466227 |
| **Fluorescein** | Alcon Nordic | 00065009265 |

| Antibody | Company | Cat No. |
|---|---|---|
| **goat anti-CD31** | R&D Systems | AF3628 |
| **rabbit anti-ERG** | ABCAM | ab92513 |
| **mouse anti-aSMA 555** | Sigma | C6198 |
| **rabbit anti-GLUT1** | Millipore | 07-1401 |
| **rabbit anti-Collagen IV** | Bio-Rad | 2150-1470 |
| **donkey anti-goat 647** | Invitrogen | A21447 |
| **donkey anti-goat Cy3** | Fisher Scientific (JacksonImmuno) | NC9056961 (705-166-147) |
| **donkey anti-rabbit 555** | Invitrogen | A32794 |
| **donkey anti-rabbit 488** | Invitrogen | A21206 |

| Solution | Compound | Amount |
|---|---|---|
| **Solution 1*** | THEED | 4-5 mL (4-5 g) |
| | Triton X-100 | 2.5 mL (2.675 g) |
| | Urea | 12.5 g |
| | $dH_2O$ | Fill to 50 mL |
| **Bleaching solution: 10% $H_2O_2$**** | 30% $H_2O_2$ | 16 mL |
| | PBS 1x | 32 mL |
| **Gelatin solution** | Gelatin | 0.3 g |
| | PBS | 10 mL |
| **PTx.2 (1L)***** | Triton X-100 | 2 mL / 2.14 g |
| | PBS 10x | 100 mL |
| | $dH_2O$ | 900 mL |
| **Permeabilization solution***** | PTx.2 | 400 mL |
| | Glycine | 11.5 g |
| | DMSO | 100 mL |
| **Blocking solution***** | PTx.2 | 42 mL |
| | Donkey Serum | 3 mL |
| | DMSO | 5 mL |
| **PTwH***** | Tween-20 | 2 mL |
| | Heparin | 1 mL of 10mg/mL stock |
| | PBS 10x | 100 mL |
| | $dH_2O$ | 900 mL |

THEED, N,N,N′,N′-Tetrakis(2-hydroxyethyl) ethylenediamine
* based on DEEP-Clear protocol (ref)
** based on EyeCi protocol (ref)
*** based on iDISCO+ protocol (ref), add 0.02% $NaN^3$ to prevent bacterial growth.

**Detailed protocol for depigmentation, immuno-staining and clearing of whole eyes**

| Step | Incubation time | Volume | Temperature | Notes |
|---|---|---|---|---|
| **Decoloration** | | | | |
| Organic- solvent: acetone | 2 hours | 2 mL | -20°C | Pre-chilled to 4°C to prevent sudden temperature difference |
| 3x wash in PBS | >5 min per wash | 2 mL | RT | With gentle agitation |
| Depigmentation In Solution-1 | 3 hours | 2 mL | 37°C | With gentle agitation |
| 3x wash in PBS | >5 min per wash | 2 mL | RT | With gentle agitation |
| Bleaching: 10% H2O2 | Variable, until sufficient bleaching | 1.5/2 mL | 37°C/55°C | With gentle agitation |
| **Immunostaining** | | | | |
| Wash in PTx.2 | 2x 1 hour | 2 mL | RT | With gentle agitation |
| Permeabilization | o/n | 2 mL | 37°C | With gentle agitation |
| Blocking | o/n | 2 mL | 37°C | With gentle agitation |
| Primary antibodies | >36 hours | 0.4 mL | 37°C | With gentle agitation |
| PTwH wash | >24 hours | 2 mL | RT | With gentle agitation, change solution 4-5x |
| Secondary antibodies | >36 hours | 0.4 mL | 37°C | With gentle agitation |
| PTwH wash | >24 hours | 2 mL | RT | With gentle agitation, change solution 4-5x |
| **Clearing** | | | | |
| Embedding in gelatin | Until solid, ~20 min | Prepare 3 mL / eye | 4°C | Process takes considerably longer, eyes should be cleaned of fibers. |
| Dehydration: 20%, 40%, 60%, 80%, 100%, 100% MeOH* | 2 hours/step | 2 mL | RT | The 100% steps are done 1h each, with gentle agitation |
| 66%DCM/33%MeOH | 3 hours | 2 mL | RT | With gentle agitation |
| 100% DCM* | 2x 15 min | 2 mL | RT | With gentle agitation |
| DBE* | >4 hours or until sufficient transparency | 2 mL | RT | Preferably longer, and with a change of DBE |

Table 1. Workflow for whole eye LSFM. RT, Room temperature; DCM, Dichloromethane; DBE, Dibenzyl Ether. *Read the safety data sheets for MeOH, DCM and DBE.

### Tutorial for Imaris based analysis of whole mouse eye vasculature using in silico dissection

An essential part of the LSFM data analysis is the in silico dissection of the different vascular compartments in the eye, as this clarifies structures by omission of others. Here we describe tools of Imaris and how they can be exploited to enhance whole eye analysis. Similar tools in open sourced/other software can naturally be used in similar manner to achieve the same goal. The *in sillico* dissection of a whole P5 mouse eye will be used as an example here.

**Before Imaris**

In our case, all the captured frames that constitute the Z-stack of the whole eye are saved individually in a folder. The individual frames need to be compiled into a single TIFF (.tif) before conversion to an imaris (.ims) file. This step can be done in the open software FIJI/ImageJ, which can be downloaded from the following domain:

https://imagej.net/software/fiji/

Once downloaded and extracted from the ZIP folder, ImageJ can be used immediately and does not require installation.

To open the individual frames in one window, go to: File > Import > Image Sequence

In the pop up window, click browse to select the folder containing the image data. Subsequently filter based on channel if applicable. A virtual stack can be used if your PC does not have good enough specs to open the whole file.

If multiple channels have been imaged, you can use the filter box to filter on images with a certain channel indication, such as 'C00', 'C01', 'C02', etc.

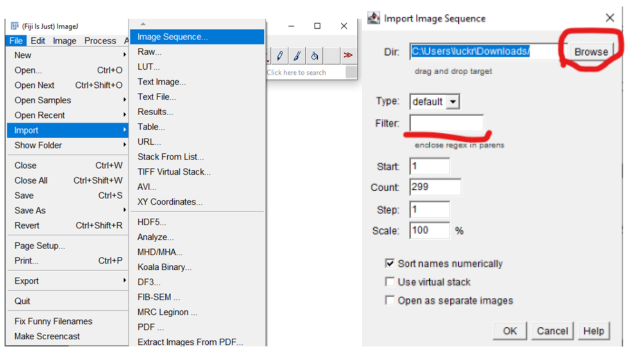


At this point some pre-processing can already be done:

- In case multiple fluorophores have been imaged and their alignment is not good enough, it is easy to adjust this in ImageJ.
    - Merge the channels by selecting: Image > Color > Merge channels
    - Subsequently check whether there is misalignment in the X,Y, or Z dimension.
    - In case of misalignment in Z:

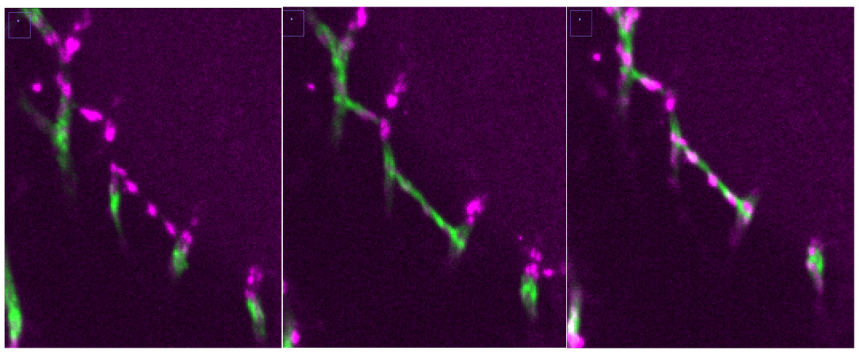

- Asses the size of the misalignment in terms of numbers of frames. Then split the channels by selecting: Image> Color > Split channels

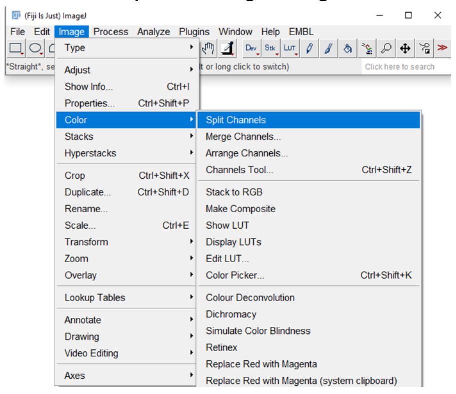


- Then open the ‘Slice Keeper’ tool by typing “keep” into the search bar, select it and ‘Run’.
- Divide the size of the estimated Z-misalignment in half and remove this number from the beginning of the stack of one channel, and from the end of the stack from the other

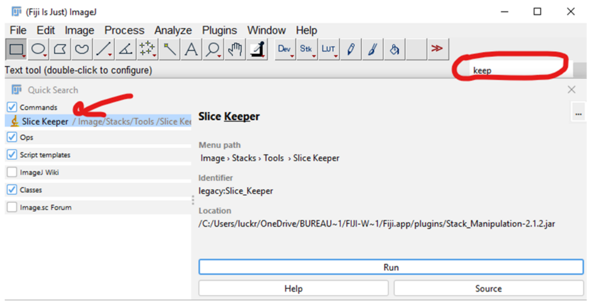


channel. Set increment to 1.

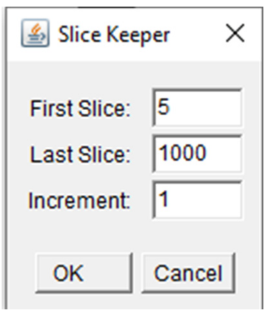


- Merge the channels by selecting: Image > Color > Merge channels

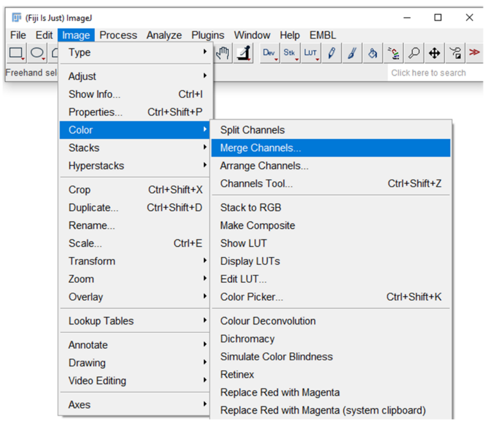


  - Check whether the channels have been properly aligned, repeat the process if not.*
- In case of misalignment in X and/or Y:
  - Assess the size of the misalignment in terms of number of pixels. Then split the channels by selecting Image > Color > Split channels
  - At this point it is useful to duplicate the channels: run the slice keeper and keep the full stacks (this goes faster than duplicating in the case of large files).
  - Select: Edit > Selection > Specify

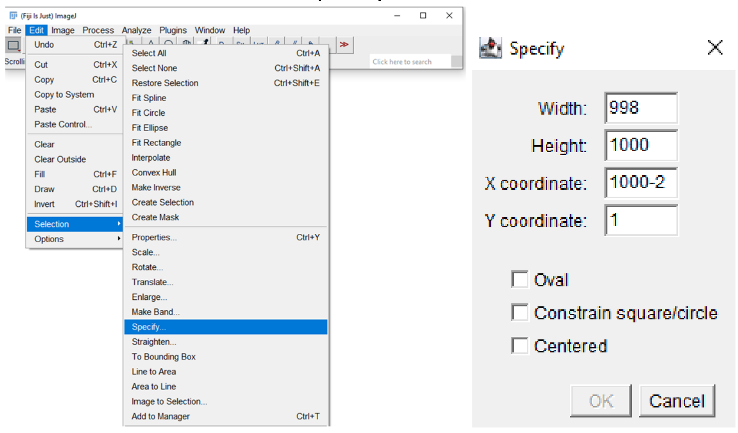


  - 
  - Specify a region of interest that is 50% of the misaligned pixels (N) smaller than the full image (in terms of width and height). Then put the coordinates on (N) for one channel and Total number of pixels (T) – N for the other channel.
  - Crop the stacks by pressing Ctrl + Shift + X
  - Merge the channels: Image > Color > Merge channels
  - Check whether the channels are properly aligned, repeat the process if not.*

- In case only a specific region needs to be analysed, it is better to crop said region out as it will save time and storage space in further analysis.
  - Select a region of interest (ROI) using one of the tools encircled below and press T.

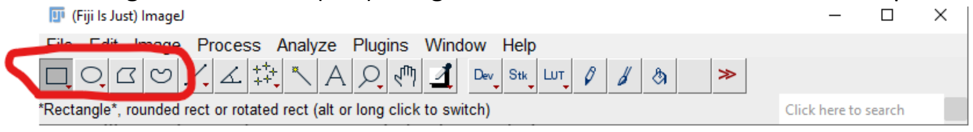


  - It is advisable to save the ROI for future reference. Go to ROI Manager (separate window) > More > Save

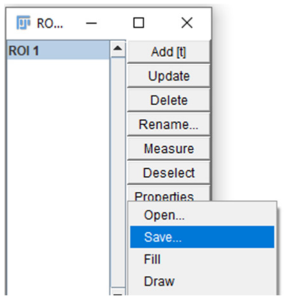


- Crop to your ROI by pressing Ctrl + Shift + X
- Extract the desired sub stack by opening ‘slice keeper’ tool (see above) and choosing the right range for your region of interest.

To improve further on our workflow, one can perform additional steps at this point, such as denoising or deconvolution algorithms.

Once all pre-processing steps are completed, the stack can be saved as a TIFF by selecting File > Save as > Tiff

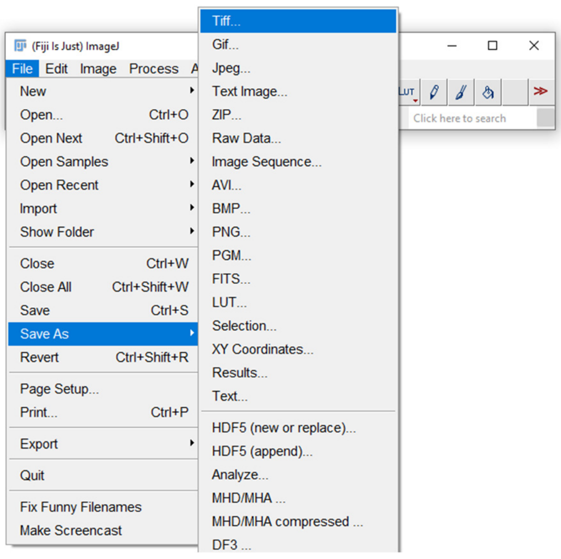


**Note: alignment of different channels helps with recognizing which structures are within proximity of each other, but does not create a ‘true’ image.*

**Dissecting the whole eye in Imaris**

To convert a TIFF to an Imaris file, open the ImarisImageConverter software. Drag the TIFF into the window, and click convert. It is useful to have it save the output in the same folder as the original TIFF.

Once converted, open Imaris, go to arena, and drag the file into the window to open it.

First, make sure the voxel size is set correctly. Press ‘Ctrl + I’ to open image properties window and set the voxel size.

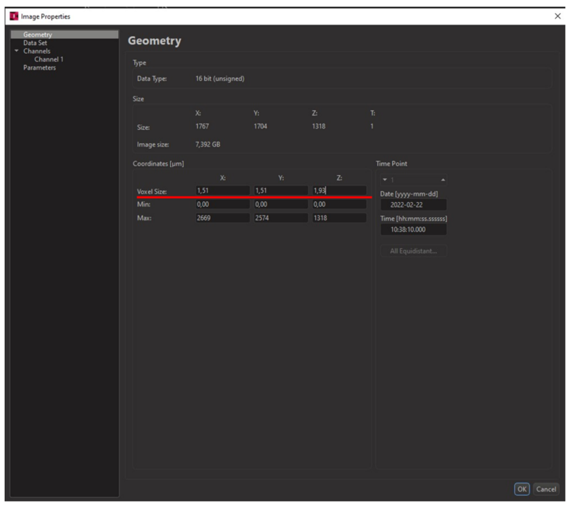


Then one can start dissecting. Select the surface tool in the top left of the screen. The choose the option 'Skip automatic creation, edit manually'.

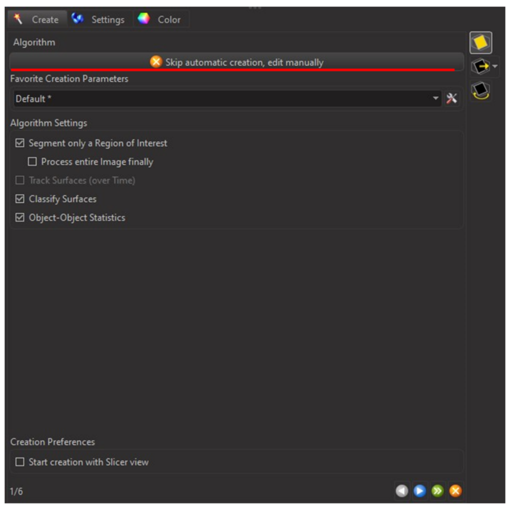


There are a few different drawing modes, we find the distance based vertex mode best to work with. With this mode a vertex will be created every set amount of distance travelled with the cursor.

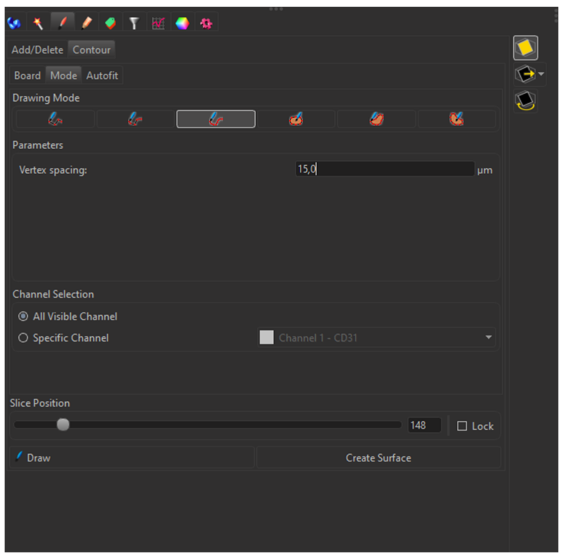

With the preferred tool selected, you can start dissecting out a desired structure. Here we will dissect out the retina, hyaloids and outer vasculature. Select the yellow plane to toggle the 'plane view' to only see the plane you are drawing on. By going to "Board" you can choose in which plane you want to draw (XY, XZ or YZ).

First we will dissect out the entire 'inside of the eye'.

Start at one end of 'the inside' and draw a circle. To see different tissue structures based on their autofluorescence, it helps to overexpose the signal. This can be done in the display adjustment window, use the right mouse button to adjust the maximum values, and the left mouse button to adjust the minimum values. (If the display adjustment window is not present, go to ' Edit' in the top left of the window and click 'Show Display Adjustment').

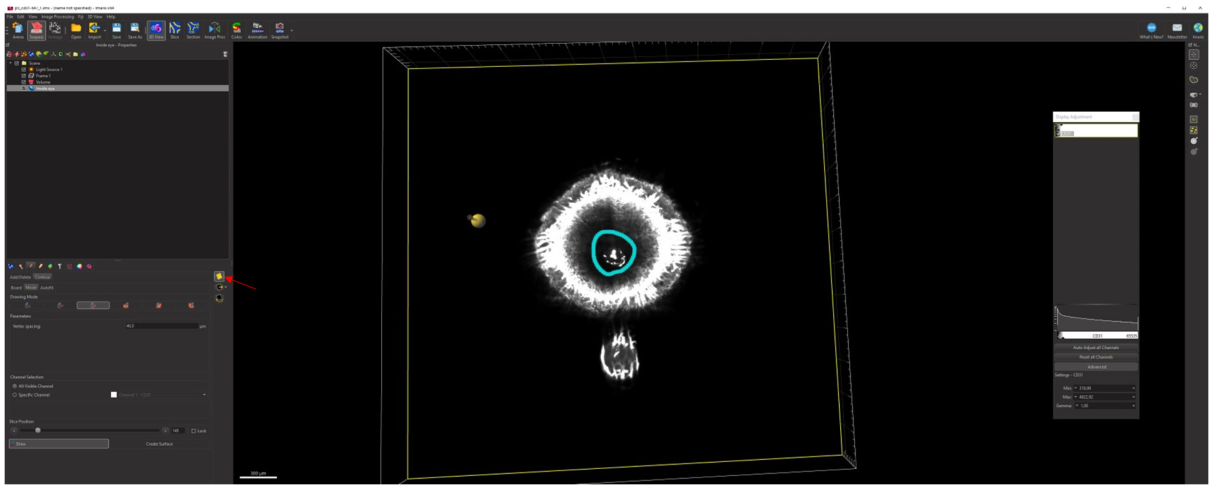

Use the yellow ball on the plane or drag in the bottom left to travel through the stack. Draw around the desired area every 50-100 slices. The more consistent a structure is in terms of shape, the less slices need to be drawn on. If a mistake is made you can press 'Ctrl + Z', or go to Board to delete a specific drawing.

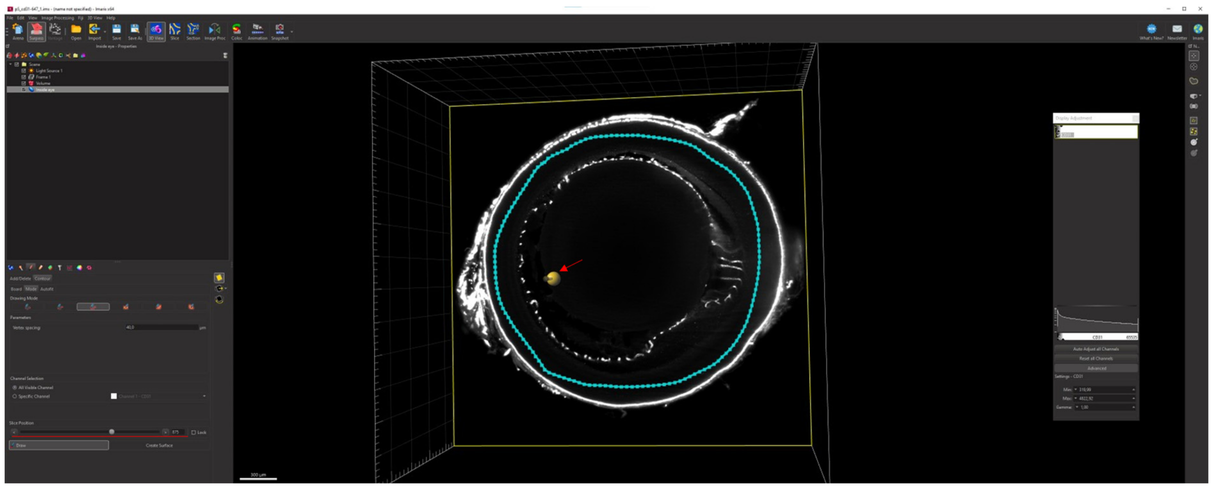

Continue this process until the end of the inside is reached (you don't have to go over the entire stack, just the structure you are isolating). Click 'Create Surface'.

Next, we create another surface object and repeat the same process, but now to dissect out the hyaloids and lens.

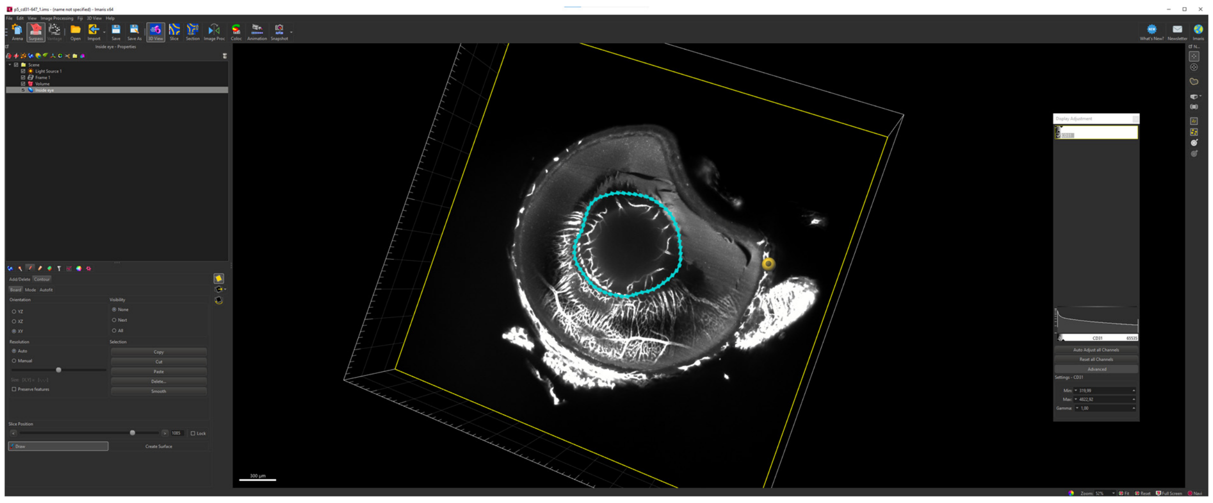

Once this is done, go back to the 'Inside eye' surface object. Click on the 'Edit' tab and subsequently 'Mask Selection' (Note: this is not the edit tab on the top of the window, but the one in the object manager). Set voxels **outside** surface to 0 and click 'Ok'.

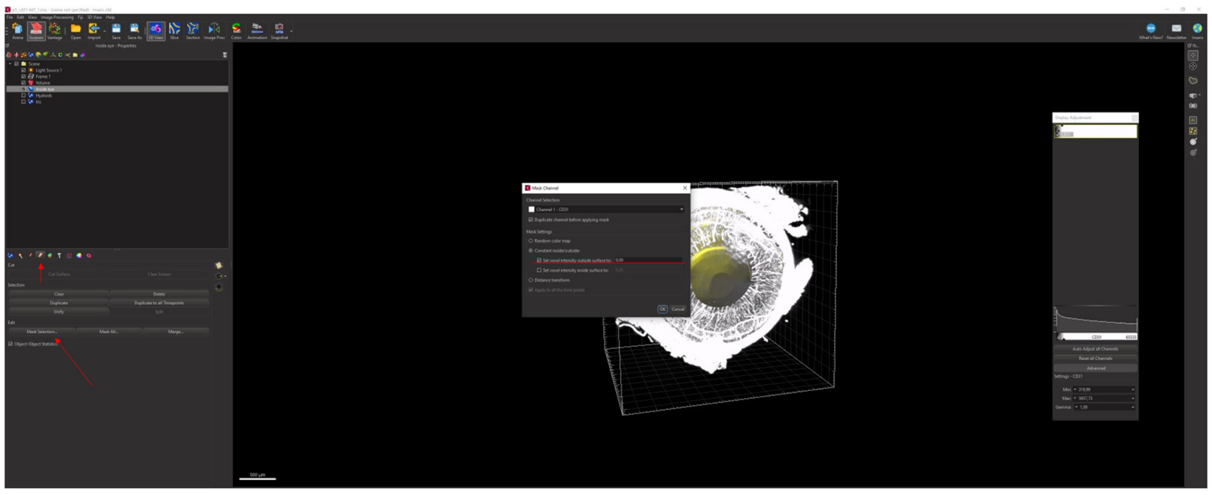

Next, do the same thing with the Hyaloids + lens surface object, but then set voxels **inside** surface to 0.

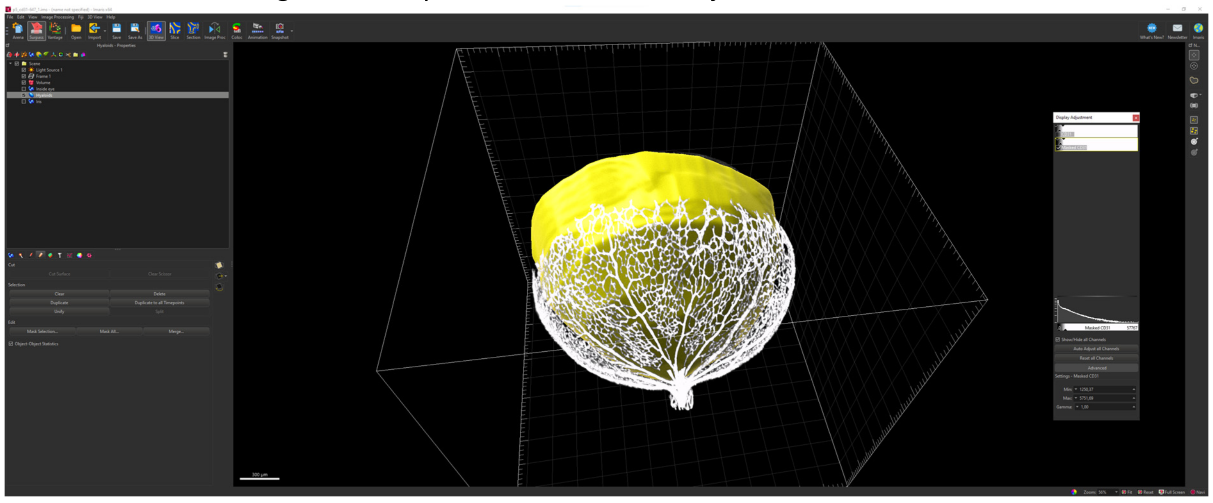

Now retina isolation is completed. It is helpful to delete channels that you do not need anymore, go to 'Edit' in the top left of the window, and click 'Delete Channels'.

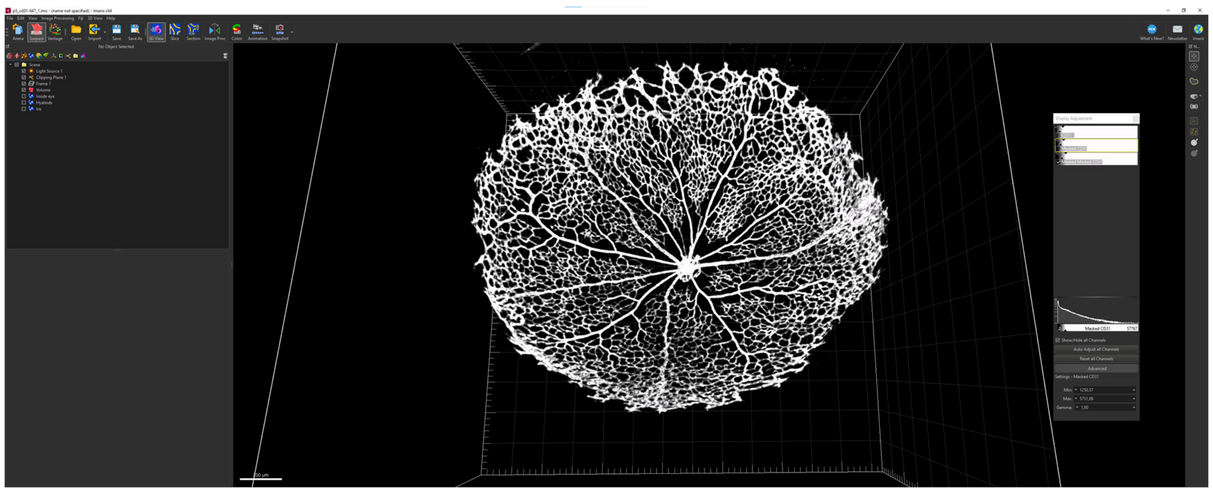

The last two steps can be reused to isolate the hyaloids and the outer vasculature.

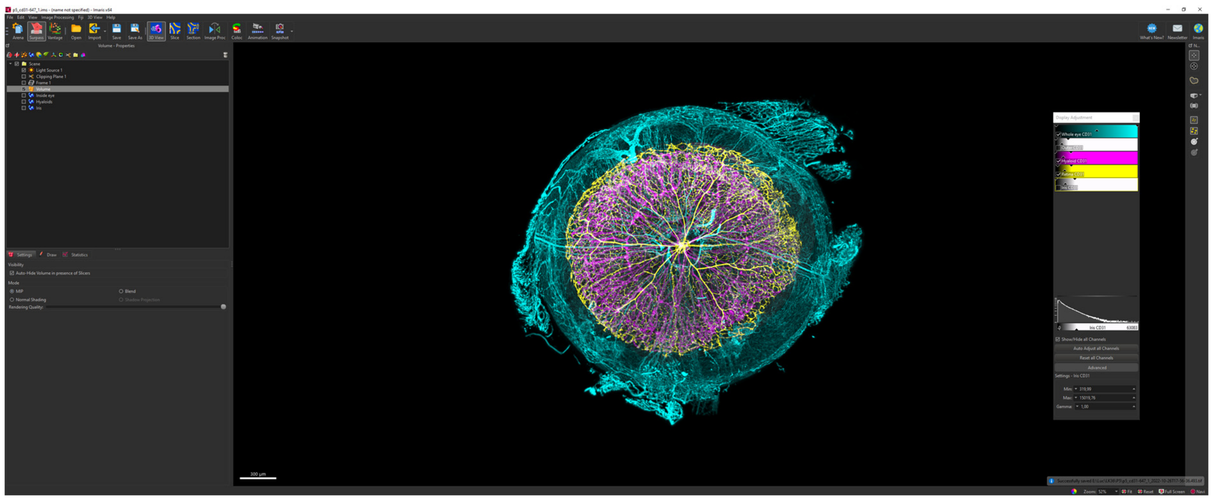

False colors, intensities of channels and toggling channels on/off can be done in the Display Adjustment window. New surface objects can be made to isolate whichever desired structure, such as the iris:

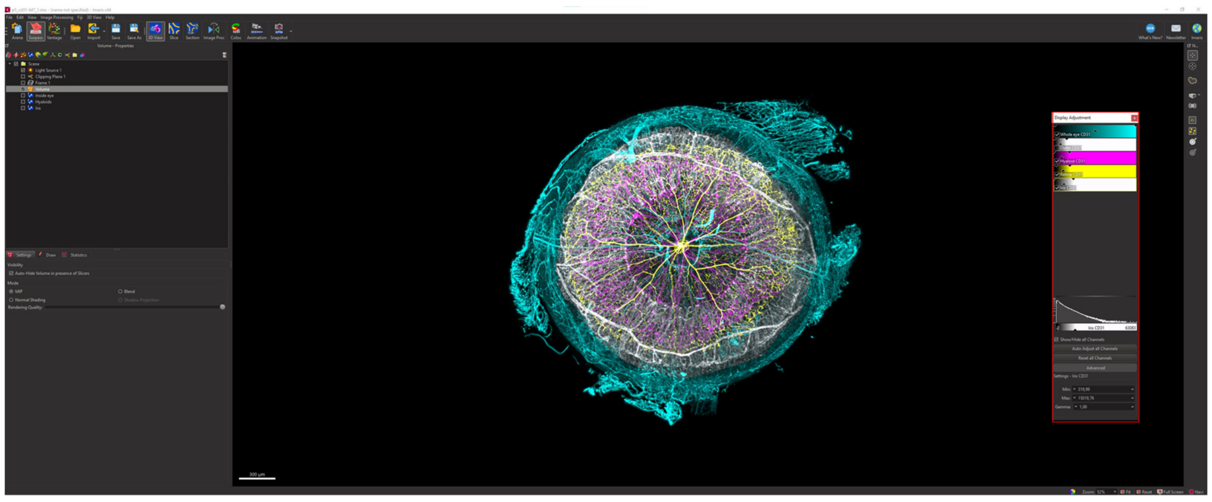

In addition to dissection through manual use of the surface tool, there are other tools that help in the visualisation of structures.

Clipping planes obscures all objects one side of the plane and can be rotated in any orientation. By using one or multiple clipping planes, you can quickly and easily put focus on a specific part of the eye and remove structures that are in the line of sight of the structure you wish too look at.

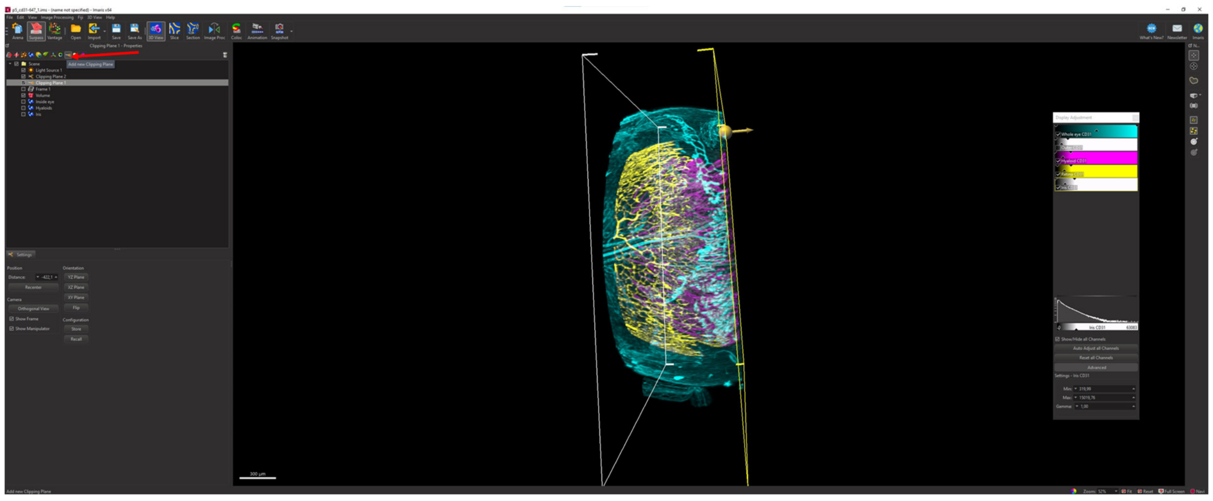

Oblique slicers allow the projection of a given section size onto a 2D plane and can also be rotated in any orientation. This can be especially useful for structures that are very thin.

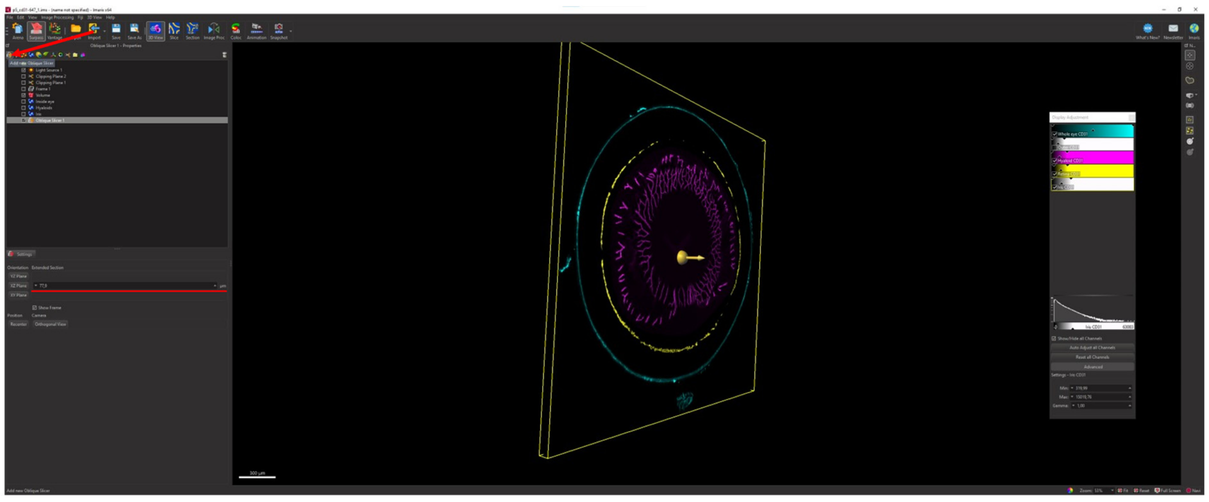
